# Gene co-expression network connectivity is an important determinant of selective constraint

**DOI:** 10.1101/078188

**Authors:** Niklas Mähler, Jing Wang, Barbara K Terebieniec, Pär K Ingvarsson, Nathaniel R Street, Torgeir R Hvidsten

**Affiliations:** Department of Chemistry, Biotechnology and Food Science, Norwegian University of Life Sciences, Ås, Norway; Umeå Plant Science Centre, Department of Plant Physiology, Umeå University, Umeå, Sweden; Umeå Plant Science Centre, Department of Ecology and Environmental Science, Umeå University, Umeå, Sweden; Centre for Integrative Genetics, Faculty of Biosciences, Norwegian University of Life Sciences, Ås, Norway; Department of Plant Biology, Swedish University of Agricultural Sciences, Uppsala, Sweden

## Abstract

While several studies have investigated general properties of the genetic architecture of natural variation in gene expression, few of these have considered natural, outbreeding populations. In parallel, systems biology has established that a general feature of biological networks is that they are scale-free, rendering them buffered against random mutations. To date, few studies have attempted examine the relationship between the selective processes acting to maintain natural variation of gene expression and the associated co-expression network structure. Here we utilised RNA-Sequencing to assay gene expression in winter buds undergoing bud flush in a natural population of *Populus tremula*, and outbreeding forest tree species. We performed expression Quantitative Trait Locus (eQTL) mapping and identified 164,290 significant eQTLs associating 6,241 unique genes (eGenes) with 147,419 unique SNPs (eSNPs). We found approximately four times as many local as distant eQTLs, with local eQTLs having significantly higher effect sizes. eQTLs were primarily located in regulatory regions of genes (UTRs or flanking regions), regardless of whether they were local or distant. We used the gene expression data to infer a co-expression network and investigated the relationship between network topology, the genetic architecture of gene expression and signatures of selection. Within the co-expression network, eGenes were underrepresented in network module cores (hubs) and overrepresented in the periphery of the network, with a negative correlation between eQTL effect size and network connectivity. We additionally found that module core genes have experienced stronger selective constraint on coding and non-coding sequence, with connectivity associated with signatures of selection. Our integrated genetics and genomics results suggest that purifying selection is the primary mechanism underlying the genetic architecture of natural variation in gene expression assayed in flushing leaf buds of *P. tremula* and that connectivity within the co-expression network is linked to the strength of purifying selection.

**Author summary:** Numerous studies have shown that many genomic polymorphisms contributing to phenotypic variation are located outside of protein coding regions, suggesting that they act by modulating gene expression. Furthermore, phenotypes are seldom explained by individual genes, but rather emerge from networks of interacting genes. The effect of regulatory variants and the interaction of genes can be described by co-expression networks, which are known to contain a small number of highly connected nodes and many more lowly connected nodes, making them robust to random mutation. While previous studies have examined the genetic architecture of gene expression variation, few were performed in natural populations with fewer still integrating the co-expression network.

We undertook a study using a natural population of European aspen (*Populus tremula*), showing that highly connected genes within the co-expression network had lower levels of polymorphism, had polymorphisms segregating at lower frequencies and with lower than average effect sizes, suggesting purifying selection acts on central components of the network. Furthermore, the most highly connected genes within co-expression network hubs were underrepresented for identified expression quantitative trait loci, suggesting that purifying selection on individual SNPs is driven by stabilising selecting on gene expression. In contrast, genes in the periphery of the network displayed signatures of relaxed selective constraint. Highly connected genes are therefore buffered against large expression modulation, providing a mechanistic link between selective pressures and network toplogy, which act in cohort to maintain the robustness at the population level of the co-expression network derived from flushing buds in *P. tremula*.

## Introduction

A central aim of biology is to understand how emergent phenotypes are encoded in the genome and how genetic variation engenders phenotypic variation within populations. While much emphasis was, and is, placed on studying the genetics of those emergent phenotypes, less attention has been paid to the genetics of the various steps along the central dogma of molecular biology (in essence, the progression of genome to RNA to protein) that underlie the emergence of a phenotype of interest. The availability of massively parallel sequencing technologies affords new possibilities for addressing biological questions, for example enabling the generation of de novo genome assemblies and population-wide resequencing data that can be used to perform genome-wide association studies (GWAS), even in species with large genomes that harbour high levels of polymorphism or that display rapid linkage disequilibrium (LD) decay [1]. The use of genome-wide resequencing data allows the discovery of, in theory, all genetic polymorphisms within an individual. These genetic markers, of which single nucleotide polymorphisms (SNPs) are currently the most commonly considered, can then be used to perform association or linkage mapping to identify the subset of polymorphisms engendering phenotypic variation among individuals.

Advances in sequencing technologies have concordantly revolutionised transcriptomics studies, particularly in non-model organisms. Following seminal work [2,3], numerous early studies in a range of species established that there is a significant heritable component underlying natural variation of gene expression levels among individuals within populations [4–16] and that this variation underlies a number of phenotypes [17–24]. Given these findings, it became apparent that gene expression values could be considered in the same way as any other quantitative phenotype and be subjected to linkage or association mapping to identify polymorphisms contributing to expression level variation among individuals [25], as first reported in [19], with the identified loci termed expression Quantitative Trait Locus (eQTL; [6]) or, less commonly, expression level polymorphisms (ELPs; [26]). eQTLs are classified as either local or distant acting depending on the physical location of the associated polymorphism in relation to the gene that the eQTL is mapped for: local eQTLs are usually defined as being located within a specified physical distance of the gene location on the same chromosome, while distant eQTLs represent polymorphisms that are located beyond that threshold distance or on another chromosome. eQTLs can be further classified as acting in *cis* or *trans*: *cis* eQTLs act in an allele-specific manner and are usually considered to be local, although long-range *cis* interactions can occur, for example when a polymorphism is located in an enhancer that is physically distant from the gene of interest; trans acting eQTLs affect both alleles of a gene and are most commonly located distant to that gene. There continues to be strong interest in eQTLs as they can identify mechanistic links between phenotype and genotype [27,28]. Importantly, the majority of polymorphisms that have been associated to phenotypes using GWAS in a wide range of species are located outside of protein coding or transcribed regions [29–32], suggesting that they influence expression rather than altering protein or transcript function.

A number of previous eQTL studies have been conducted using plant species including *Arabidopsis thaliana* [33–38], *Zea mays* [39–41] and *Oryza sativa* [42,43], and in forest tree species [7,44–46]. These, together with studies in other eukaryotic systems, have yielded generalities concerning the genetic architecture of gene expression variation, including that a greater number of local eQTLs are typically identified and that these individually explain a larger proportion of gene expression variance than do distant eQTLs [34,47–50]. Much of the previous work was conducted using controlled, and for tree species in particular inter-specific, crosses, comparisons of accessions or was performed in non-natural systems and it is not clear how generally applicable their conclusions are for natural populations of unrelated individuals. Few studies have considered whether observed, heritable variation is adaptive [51,52] or whether signatures of population differentiation are observed at the transcriptome level [52–57], and it is not yet clear the extent to which selection acting on gene expression underlies adaptive phenotypic trait variation [52].

Systems biology has greatly improved our understanding of the shared regulation of genes, revealing the topological properties of transcriptional co-expression networks. A salient feature of the topology of co-expression networks is that they are scale-free, having few highly connected nodes (genes) and many nodes with few connections. This property imparts an inherent ability to buffer against single mutations of large negative effect as random mutation of an expression pattern (*i.e*. an eQTL) or coding sequence will more often affect a network node of low connectivity [58-60]. Although there have been a number of eQTL studies performed, the context of eQTLs within the co-expression network and how this relates to patterns of selection have not been a focus.

Species in the *Populus* genus have been established as a powerful model system for forest tree genomics due to their relatively small genome, rapid growth, propensity for clonal propagation and ease of genetic transformation [61]. *P. tremula* (European aspen) has many features that render it a particularly useful model for population genetics and speciation studies [62,63], studies of which are facilitated by availability of a draft *de novo* genome assembly [64] and population resequencing data [65]. In this study, we aimed to determine the evolutionary forces that maintain the genetic variation of gene expression within the context of the corresponding co-expression network using a natural collection of *P. tremula*. Specifically, we wished to test whether co-expressed sets of genes are enriched for specific biological functions, whether network topology influences gene expression and sequence evolution of the constituent genes, and how selection interacts with network topology to affect the patterns of genetic variation within populations. To address these questions we generated population-wide RNA-Seq data in, assaying gene expression in winter buds at the point of bud break. We performed eQTL mapping and constructed a co-expression network, which was scale free and modular, with highly connected genes in the module cores being under-represented for eQTLs and with eQTL effect size being negatively correlated with gene connectivity. Patterns of polymorphism and divergence within genes in module cores imply that they are likely experiencing stronger selective constraint relative to genes in the network periphery. Our results suggest that purifying selection plays an important role in buffering the transcriptional network against large perturbations and that natural variation in gene expression is more prevalent in genes of low network connectivity as a result of relaxed selective constraint.

## Results

We utilised the northern common garden (located at 63.9° N, near Umeå, Sweden) of the Swedish Aspen (SwAsp) collection [66], which comprises 116 *P. tremula* genotypes sampled from twelve geographic locations spanning the species distribution range in Sweden (56.2° to 66.4° N, Fig 1A). The SwAsp collection was previously shown to contain abundant genetic variation, with low linkage disequilibrium (LD) [63,67] and minimal population structure [68]. Based on whole genome re-sequencing data aligned to a *de novo* assembly of the *P. tremula* genome (available at PopGenIE.org; [64]), we called 4,509,654 SNPs after stringent filtering, which we utilised to perform eQTL mapping. To examine the genetic architecture of natural variation in gene expression within the SwAsp collection, we generated paired-end RNA-Seq expression data from winter buds at the point of spring bud break/flush for 219 individuals (clonal replicates), representing 86 genotypes.

**Fig 1.**
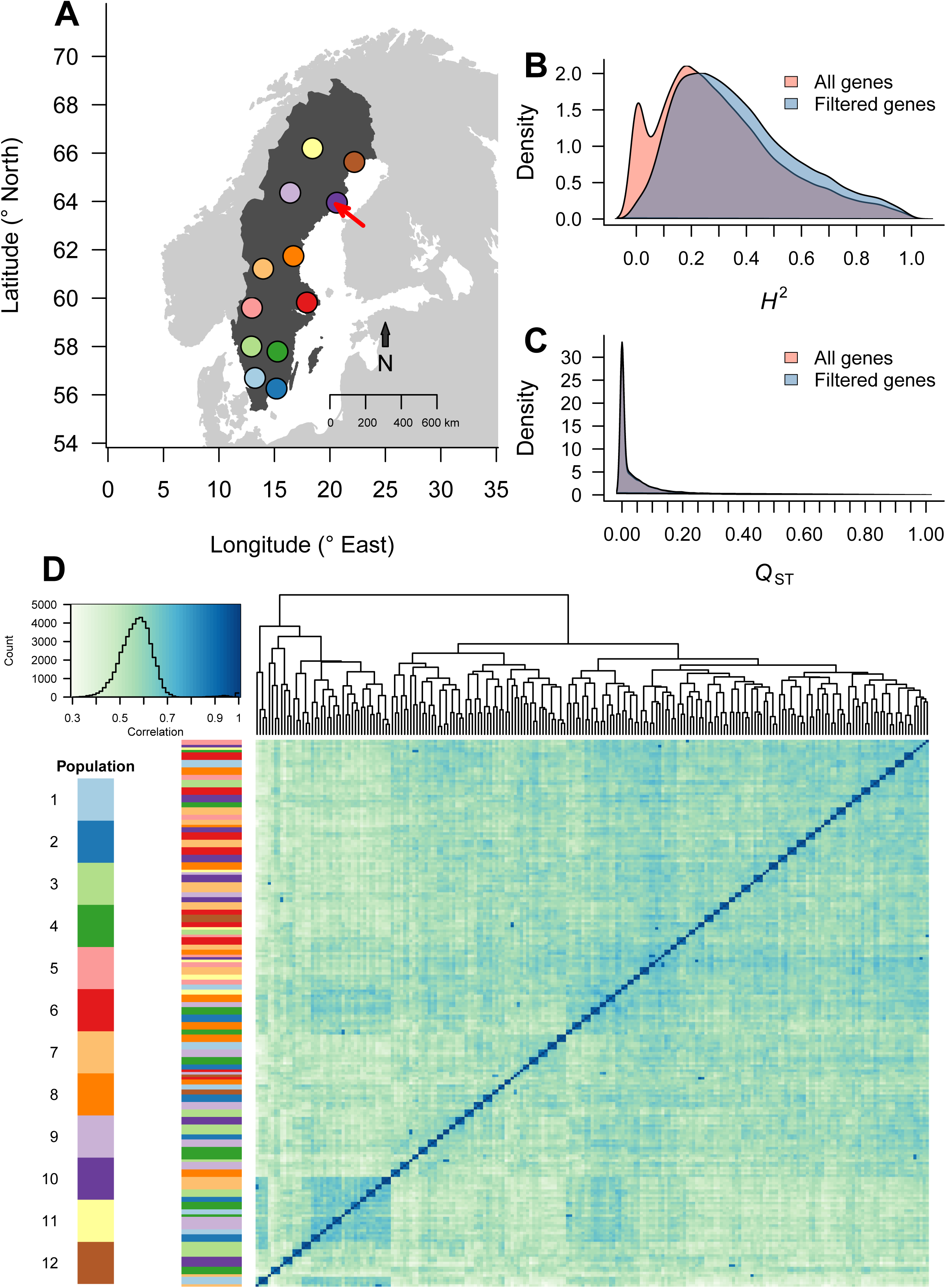
Population location and gene expression overview. (A) Map of the original locations of the SwAsp populations. The red arrow points to the location of the common garden used in this study. (B) Distribution of gene expression heritability for all genes and for the subset of genes after filtering to remove uninformative expression. (C) Distribution of gene expression *Q*_ST_ for all genes and for the subset of genes after filtering to remove uninformative expression. (D) Sample clustering based on all samples, including biological replicates. The heatmap represents the sample correlation matrix based on the 500 genes with the highest expression variance using gene expression data prior to hidden confounder removal. Darker colour indicates higher correlation. The coloured bar represents the populations the samples belong to. The small clusters on the diagonal correspond to biological replicates of each genotype.

### Population level gene expression similarity

We first examined the distribution of broad-sense heritability (*H^2^*) and population differentiation (*Q*_ST_) (Fig 1B,C respectively) for all expressed, annotated genes. *H^2^* ranged from 0.0 to 1.0 with a mean (± s.d) of 0.30 (0.22) and with 5,924 genes (17%) having H^2^ > 0.5. Permutation testing showed that 21,219 genes had significantly higher heritability than expected by chance (p < 0.01). There was a weak positive correlation between *H^2^* and median expression level (Pearson r = 0.09; df = 32,767; p < 2.2×10^-16^), and a relatively strong positive correlation to expression variance (Pearson r = 0.43; df = 32,767; p < 2.2×10^-16^). *Q*_ST_ ranged from 0.0 to 1.0 with a mean (± s.d) of 0.06 (0.12) and had a weak negative correlation with expression variance (Pearson r = -0.02; df = 29,670; p < 4.5×10^-4^) and a positive correlation with median expression (Pearson r = 0.18; df = 29,670; p < 2.2 ×10^-16^). These findings are similar to those reported for a number of species [5–17,53], suggesting that the expression of a large proportion of genes is under substantial genetic control and that the expression of highly expressed genes is under generally tighter genetic control than genes with lower expression.

While a number of previous studies have identified evidence for heritability of gene expression [52,69,70], the relationship between expression variation and population structure has been explored less. Our previous work has established that there is minimal population structure at the genetic level in the SwAsp collection [68]. To examine whether population structure was apparent on the basis of expression variation among genotypes, we performed hierarchical clustering of all individuals (Fig 1D) or genotypes (S1 Fig). While some evidence of clustering among genotypes was apparent (regions of blue in Fig 1D, S1 Fig), genotypes (or individuals) did not cluster according to population of origin (as indicated by the y axis color bars), suggesting that the observed clustering does not result from population structure. We tested whether the clustering could be used to predict the population of origin for genotypes by cutting the dendrogram to produce 12 clusters that were used as the response in a multinomial logistic regression. The mean accuracy for a 10-fold cross validation was extremely low (0.09). We additionally performed permutation tests, which showed that the mean *Q*_ST_ for all genes was significantly lower than expected by chance (p < 0.001).

To identify whether genes with the highest *H*^2^ and *Q*_ST_were enriched for characteristic biological functional signatures we selected the 500 genes with the highest *H*^2^ (0.88-1.0, 0.93 ± 0.03) and 500 genes with the highest *Q*_ST_ (0.54-1.0, 0.71 ± 0.13) and subjected these to GO enrichment analysis (see S1 File for all results). Genes with high *H*^2^ were enriched for categories including protein phosphorylation (GO:0006468; p = 3.7×10^-6^), while high Q_ST_ genes were enriched in terms including translation (GO:0006412; p = 4.2x10^-20^) and gene expression (GO:0010467; p = 4.6×10^-12^). Likewise, we considered the genes with the lowest values, which revealed enrichment of terms including cell wall modification (GO:0042545; p = 2.6×10^-7^) for the 2,289 genes with an *H*^2^ of zero and enrichment of terms including amino acid activation (GO:0043038; p = 0.0012) among the 11,895 genes with a *Q*_ST_ of zero.

We performed a regression analysis to ascertain whether a set of geographic (latitude, longitude, elevation), climatic (temperature, precipitation) or other (time since sample collection) factors significantly explained the global patterns of gene expression similarity among genotypes (S2 Fig), as identified by performing a PCA of the expression data. None of the gene expression principal components (PCs) were significantly explained by these environmental factors, with the only significant results found between PCs 2, 5 and 7 and the number of hours from collecting branches from the field until bud samples were collected in the greenhouse, which explained 6.6%, 3.2%, and total 2.1% expression variance, respectively.

We subsequently filtered expression values to remove unexpressed genes and uninformative expression profiles with low variance, as these are uninformative for association mapping or for co-expression analyses. Of 35,154 annotated genes, 20,835 were expressed in all samples, including biological replicates, and 23,183 were expressed in all genotypes when considering genotype means. Filtering to remove uninformative expression retained 22,306 genes, with the 12,848 removed genes representing those that were either not expressed in our bud samples (6,736 genes with median expression of zero of which 2,385 had no detectable expression at all), or that were weakly expressed (1,762 genes with variance < 0.05 and median expression < 2), together with genes that had stable expression among genotypes (4,350 genes with expression variance < 0.05 and median expression >= 2). The latter potentially represent genes with canalised gene expression. Analysis of this set of stably expressed genes identified enrichment for GO categories including protein transport (GO:0015031, p = 6.8×10^-11^) and protein localisation (GO:0008104, p = 2.2×10^-10^). In contrast, the 500 genes with the highest variance across all samples were enriched for GO categories related to protein phosphorylation (GO:0006468, p < 10^-6^), chitin metabolic process (GO:0006030, p < 10^-4^), and cell wall macromolecule catabolic process (GO:0016998, p < 10^-4^). Comparing the variance of these 500 genes with mean F_ST_ calculated using SNPs within those genes revealed no apparent relationship, suggesting that these patterns were not the result of population structure.

A recent reanalysis [71] of two existing datasets assaying gene expression among natural accessions of *A. thaliana* [72,73] observed that thousands of genes displayed clear present/absent expression among accessions. In contrast, when filtering our data using a similar approach, we did not find any genes displaying this pattern of expression variation (S3 Fig), an observation that we also confirmed in an independent *P. tremula* dataset [74], albeit containing substantially fewer genotypes.

### eQTL mapping

To explore the genetic architecture of gene expression variation among genotypes, we performed eQTL mapping, defining an eQTL as a significant association between a SNP (termed an eSNP) and the expression of a gene (termed an eGene). Furthermore, we classified an eQTL as *local* if the eSNP was located on the same chromosome and not more than 100 kbp from the associated eGene, and as *distant* otherwise. Our threshold distance for local/distant classification was empirically determined based on the distribution of distances between eSNPs and their associated genes and the assumption that most detectable eQTLs located within one chromosome were local (S4 Fig). We did not consider whether eQTLs acted in *cis* or *trans*. In common with other studies [75–77] we removed hidden confounders in the expression data prior to mapping eQTLs by removing variance attributable to the first nine PCs of the expression data, removal of which maximised the number of eQTLs identified (S5 Fig). After removing these hidden confounders, we repeated the gene expression clustering analysis and observed that the previous sample clustering was no longer apparent (S6 Fig).

In total we identified 164,290 eQTLs at a 5% empirical FDR: 131,103 local and 33,187 distant. These eQTLs represented pairwise associations between 6,241 unique genes (eGenes; 28% of genes considered) and 147,419 unique SNPs (eSNPs), with a mean of 21.0 local and 5.3 distant eSNPs per eGene, respectively. 4,091 genes had only local eQTLs, 1,050 had only distant eQTLS while 1,100 had both. Local eSNPs explained significantly more of the variance than distant eSNPs (local mean adjusted %VE = 51, distant mean adjusted %VE = 47, Mann-Whitney p < 2.2×10^-16^, Fig 2A) and also had higher statistical significance (Mann-Whitney p = 6.9×10^-12^, Fig 2E). As expected there was a clear tendency for a local eSNP to be located proximal to the transcription start site (TSS) or the stop codon (S10 Fig). eGenes had 229 significantly higher heritability than non eGenes (median heritability difference was 0.16, permutation 230 test p < 0.0001) (Fig 2B), with this trend being slightly higher for local than distant eQTLs (S25A Fig). There was also an expected, positive correlation between the maximum %VE of the eSNPs associated with an eGene and gene expression *H*^2^ (Pearson r = 0.47, df = 6,232, p < 2.2×10^-16^). These patterns are broadly similar to those reported in a number of previous studies [34,47–50], although the ratio of local-to-distant eQTLs differs among studies and is highly influenced by sample size. Before hidden confounder removal, eGenes had marginally higher mean expression than non-eGenes (mean expression 3.5 and 3.3, respectively; permutation test p-value < 0.0001). There were no significant differences after hidden confounder removal, regardless of whether the eQTL was local or distant. eGenes with at least one local eQTL were enriched for GO categories related to tRNA metabolic process (GO:0006399, p = 1.5×10^-5^), ncRNA metabolic process (GO:0034660, p = 2.6×10^-5^) and organonitrogen compound biosynthetic process (GO:1901566, p = 2.2×10^-5^), among others, while eGenes with at least one distant eQTL were enriched for categories including protein phosphorylation (GO:0006468, p = 0.0064; see S1 File for all results).

**Fig 2.**
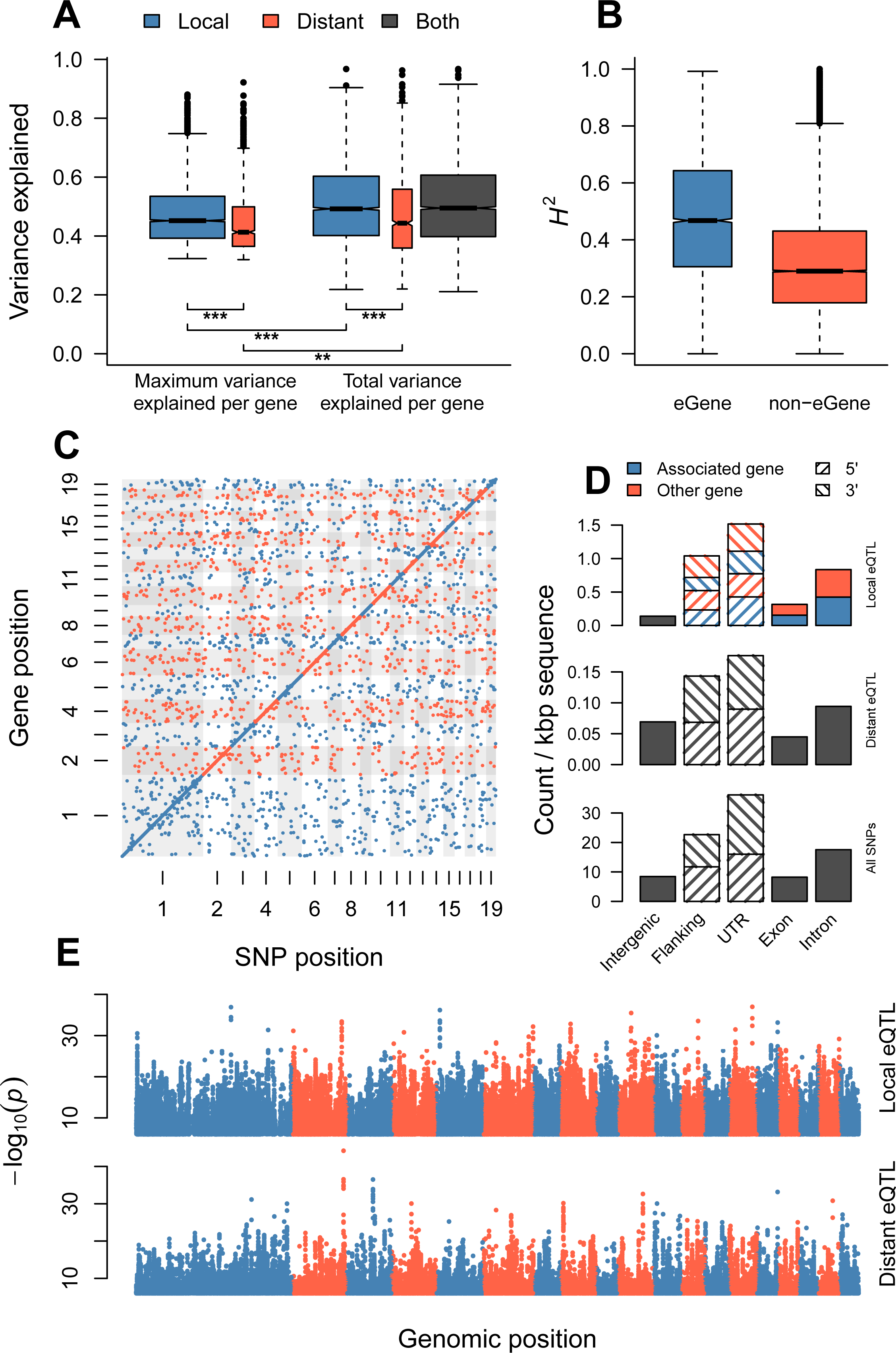
eQTL overview. (A) Expression variance explained (*R^2^*) for local and distant eQTLs. Box plots show the maximum variance explained by a single eQTL for each gene and the total variance explained by all eQTLs for each gene. The widths of the boxes are proportional to the number of genes represented. The pairwise significiance of a Mann-Whitney test is indicated by asterisks: ***p < 0.001, **p < 0.01. (B) Broad sense heritability distributions for eGenes and non-eGenes. (C) Scatter plot showing the positions of all significant eQTLs in this study. No evidence of eQTL hotspots can be observed. Numbers indicate chromosome. (D) Genomic context of local and distant eSNPs. Context categories are normalized for feature length. When an eSNP overlapped with features on both strands, both of them were counted. For both local and distant eQTLs the features are based on the gene that is closest to the eSNP, and furthermore, for local eQTLs, the features are divided into whether the eSNP is located in or near the same gene that it is associated to (“associated gene”) or not (“other gene”). Flanking regions represent 2 kbp upstream and downstream from the gene. (E) Manhattan plots for local eQTLs (upper) and distant eQTLs (lower). Each point represents an eQTL.

In contrast to a number of previous studies [19,34,36,47,50], we did not find evidence for any distantly acting hotspots (Fig 2C, S7 Fig), which represent loci where large numbers of trans-acting variants are co-located. Although the removal of hidden confounders has been shown to improve the signal:noise ratio for eQTL mapping [75–77], it is possible that the process may remove the signature of large effect distantly-acting hotspots. We performed eQTL mapping before hidden confounder removal and observed one hotspot representing 12 eSNPs assocaiated with the expression of 278 genes that was not present after removing hidden confounders (S8 Fig). The 12 eSNPs are located in close physical proximity and are therefore likely linked. In our data, the vast majority of eSNPs were associated with a single eGene (132,258 eSNPs) with a maximum of six eGenes associated with a single eSNP (S9A Fig). In contrast, only 1,248 of the 6,241 eGenes were associated with a single eSNP, with eGenes associated with up to 1,547 eSNPs (S9B Fig). In cases where eSNPs associated with the expression variation of a single eGene are physically close together, these eSNPs may be identified due to linkage rather than all being causative. To account for this we fitted linear models between the expression of each eGene and all the significant eSNPs for that gene, both local and distant. The use of a linear model masks eSNPs that contain identical/redundant information and thus effectively identifies haplotype blocks present in all individuals (which we refer to as ‘unique eSNPs’), while also producing a measure of how well the combination of eSNPs explains the expression of the corresponding eGene (in terms of percentage variance explained, %VE). Of the 4,993 eGenes associated with more than one eSNP, 4,703 were also associated to more than one unique eSNP, of which 4,210 genes were associated with at least one local eSNP and 1,203 were associated with at least one distant eSNP. The adjusted %VE for the combination of eQTLs was, in general, higher (mean %VE 51.1) than for single eSNPs (mean %VE 44.3).

We next considered the genomic context of eSNPs, which was determined by intersecting eSNP positions with gene annotations. After normalising for feature length, the majority of local eSNPs were located within untranslated regions (UTRs) and up- or down-stream (regulatory) regions of genes, with distinctly lower representation within exons than introns (Fig 2D). The genomic context distribution of local and distant eSNPs was largely similar, although there were distinctly more eSNPs located within intergenic regions for distant eQTLs. These distributions patterns are consistent with previous findings in natural populations of humans [76,78], *Drosophila* [79] and *Capsella grandiflora* [27]. A local eSNP/eQTL can be located within the region of the associated eGene (5’/3’ 2 kbp flanking, 5’/3’ UTR, exon, intron), within the region of a gene other than the associated eGene or within an intergenic region. We found that approximately half of the local eSNPs were located within the region of the eGene itself and half within another gene, with relatively few local eSNPs located in intergenic regions. When the eSNP was located within the gene region of the associated eGene there was a clear tendency for that eSNP to be located proximal to the transcription start site (TSS) or the stop codon (S10A Fig). This patterns was not present in the cases where the eSNP was located in the adjacent or another gene (S10B,D Fig), even after accounting for strand (S10C,D Fig). In these cases there was also a lower tendency for the eSNP to be located within the gene body, with a generally higher presence of eSNPs in the flanking gene regions. Given this pattern, we therefore examined the expression correlation of the eGene to the gene in which the eSNPs was located, contrasting this to pairs of non-eGenes and pairs where the eSNP and located within the eGene and the adjacent gene (S11 Fig). For those cases where the eSNP was not located within the eGene, we observed a higher expression correlation between the eGene and the gene in which the eSNP was located, potentially indicating that the eSNP induces a local and more general influence on expression. As UTRs are currently not well annotated in *P. tremula* it should be noted that many SNPs currently classified as being located in flanking regions may actually reside within UTRs. The global distribution of genomic contexts for all investigated SNPs (regardless of whether they were eSNPs or not) was similar to that of both local and distant eSNPs, suggesting no notable ascertainment bias for eSNPs.

### Co-expression network

Systems biology studies typically consider datasets assaying gene expression throughout development, among tissue types or in response to abiotic, biotic or genetic perturbations. A characteristic and salient feature of the resultant co-expression networks is that they are scale-free[80]. To determine whether the co-expression network representing expression variation among individuals within our natural population displayed the same properties, we used genotype mean gene expression values, after removal of hidden confounders, to calculate a co-expression network. In common with other biological networks, the network was scale-free (R^2^ = 0.97), suggesting that the genetic polymorphisms underlying the observed expression variance induce similar co-expression structures to those observed in previous systems biology studies. We compared the correlation and variance properties of our dataset to that of the *P. tremula* expression atlas (exAtlas; [64]), which represents different tissues collected from a single genotype. The correlation distribution for the exAtlas samples was much wider (mean correlation 0.01 ± s.d. 0.36) than that of our population expression data (mean correlation 0.00 ± s.d. 0.12; Kolmogorov-Smirnov D = 0.14, p < 2.2×10^-16^). The expression variance for the SwAsp expression data was also significantly lower than in the exAtlas data (Wilcoxon signed rank test, V = 18274602, p < 2.2 ×10^-16^; Fig 3A).

**Fig 3.**
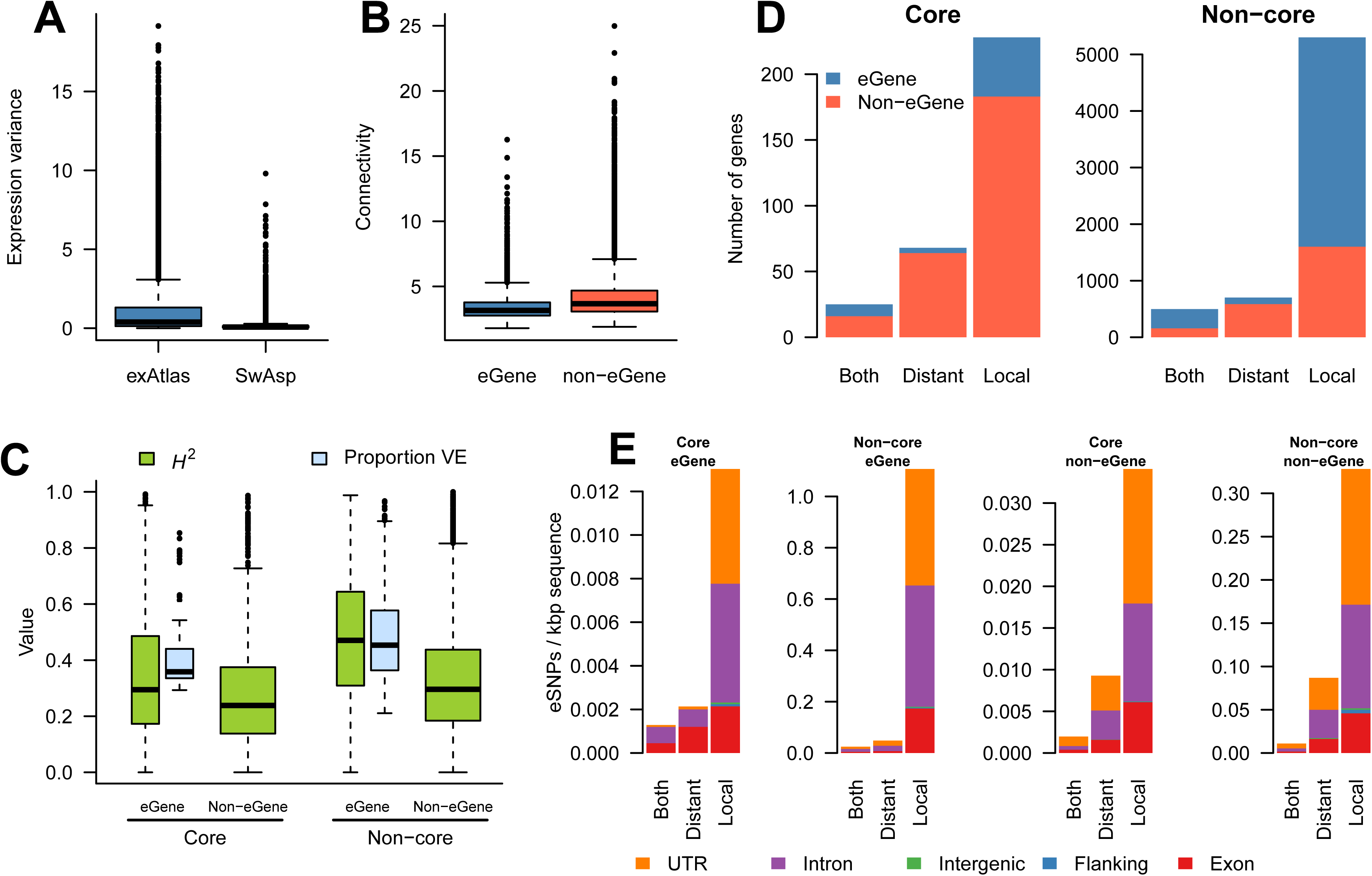
Co-expression characteristics of eGenes and eSNPs. (A) Gene expression variance distribution for all genes in the SwAsp data (before removal of hidden confounders) and the exAtlas data. (B) Distribution of co-expression connectivity for eGenes and non-eGenes. (C) Distributions of the proportion of total variance explained and heritability for eGenes and non-eGenes divided into core and non-core genes. (D) Genes having distantly, locally, or both distantly and locally acting eSNPs located within 2 kbp (or inside) the gene divided into core and non-core genes. (E) Genomic context of distantly, locally, or both distantly and locally acting eSNPs located within 2 kbp of an eGene divided into core and non-core eGenes. The eSNP counts are normalised for total feature length.

Clustering analysis of the co-expression network identified 38 co-expression modules (S12 Fig) containing a total of 20,686 genes (min=86 genes, max=1591 genes). These were enriched for a number of Gene Ontology (GO) categories including translation (modules 9, 10, and 14), photosynthesis (module 22) and oxidation-reduction process (module 29; for all results see S1 File). Despite the narrow distribution of correlation values, the modules were reasonably well defined, as indicated by the normalised connectivity difference (*k*_diff_), i.e. the difference between intra- and inter-modular connectivity. All modules exhibited a positive mean *k*_diff_), with only 157 genes (0.7%) having a negative *k*_diff_. This was in stark contrast to genes assigned to the ‘junk’ module (i.e. all genes not assigned to any well-defined module), where there were 480 genes with negative *k*_diff_ (29%).

### eGenes are under-represented in network module cores

We examined the relationship of eGenes and eQTLs to network connectivity, determining significance of these results using permutations tests shuffling gene assignments while maintaining the network structure (*i.e*. node gene IDs were shuffled while edges remained constant). In general, eGenes had lower connectivity and betweenness centrality than non-eGenes, but the effect was minimal (mean difference of 0.79 and 8.9×10^-5^, respectively; permutation test, p < 0.0001 for both, Fig 3B). Moreover, genes with a positive *k*_diff_ were significantly under-represented for eGenes (permutation test, p < 0.0001). We defined the core of each module as the 10% of genes in a module with the highest normalised *k*_diff_ while also having an intra-modular connectivity >1. Using this definition, all 38 modules contained at least one core gene, with the percentage of core genes ranging from 2-10% (S2 File, S12 Fig). Before removal of hidden confounders, core genes had both higher mean expression and variance than non-core genes (difference of 1.1 and 0.16, respectively; permutation test, p < 0.0001). However, the co-expression network was inferred from the expression data after the removal of hidden confounds, which removed these difference (no difference in mean expression: permutation test p = 0.39, and marginal difference in variance: mean variance of 0.12 and 0.11, permutation test p = 0.0184, S13 Fig). Additionally, there was a weak negative relationship between network connectivity and gene expression variance in the network as a whole (Pearson r = - 0.08; df = 22,304; p < 2.2×10^-16^; S14 Fig) and eGenes of the highest connectivity had lower effect size (based on the maximum effect size across all eQTLs associated to each eGene, Pearson r = - 0.15; df = 6,239; p < 2.2×10^-16^, S15). Among the module cores, 28 contained at least one eGene, with 25 module cores being significantly under-represented for eGenes (permutation test, p < 0.05; Fig 3D). This further emphasised that, in general, eGenes were not central in the network. There was at least one eSNP located within the region of a core gene (5’/3’ flanking, 5’/3’ UTR, exon, intron) for 32 of the 38 modules, with module cores also being under-represented for eSNPs (permutation test, p < 0.0001). For a number of metrics, module 23 had notable differences to the general pattern: It contained few core genes, all of which were eGenes; the core genes had high variance compared to non-core genes; the module in general was enriched for eGenes.

We tested whether the periphery of the network was over-represented for eGenes. Sixty-four of 145 peripheral genes were eGenes, representing a significant enrichment (permutation test, p < 0.0001). On the other hand, network module cores were enriched for both transcription factors (permutation test, p < 0.0001), which had higher connectivity than non-transcription factors (permutation test, p <0.0001), and phylogenetically conserved genes (permutation test, p < 0.0001), defined as genes with orthologs in *P. tremuloides, P. trichocarpa* and *A. thaliana* (20,318 genes; [81]). Furthermore, *P. tremula-*specific genes, *i.e*. genes without orthologs in *P. tremuloides, P. trichocarpa* or *A. thaliana* (1,614 genes), were slightly under-represented in network module cores (permutation test, p = 0.009) while being slightly over-represented among eGenes (permutation test, p = 0.0076).

In addition to eGenes having generally lower connectivity, there was also a negative relationship between eQTL effect size and co-expression connectivity for both local (Pearson r = -0.15; df = 5,189; p < 2.2×10^-16^) and distant eQTL (Pearson r = -0.12; df = 2,148; p < 6.2×10^-8^). The *H^2^* of eGenes within the core was lower than for eGenes outside the core (mean difference of 0.10, permutation test p < 0.0001, Fig 3C) and *H^2^* correlated negatively with connectivity in the network as a whole (Pearson r = -0.30; df = 22,304; p < 2.2×10^-16^).

We examined the distribution of the mode of action (Fig 3D) and genomic context (Fig 3E) of eSNPs within the network. There were distinctly more distantly acting eSNPs within the core than the non-core (Fig 3D) and, of these, there were more distantly acting eSNPs located within exons of core eGenes compared to core non-eGenes, which had higher representation of distantly acting eSNPs located in UTRs (Fig 3E). The genomic context distribution of local acting eSNPs was similar in all cases (core/non-core and eGene/non-eGene; Fig 3E), however it is clearly apparent that non-core eGenes contained by far the greatest density of eSNPs.

### Paralogs with diverged expression are more likely to be eGenes

The Salicaceae lineage underwent a relatively recent (58 million years ago) whole-genome duplication (WGD) shared by all member species and that remains represented by a large number of paralogous gene pairs in the genomes of *Populus* species [82]. If many of these duplicated genes are functionally redundant or in the process of diverging, one would expect them to be overrepresented for eGenes as sub- or neo-functionalisation requires derived SNPs to drive expression or coding divergence. To test for evidence of this we considered paralogous pairs of genes derived from the WGD event. In *P. tremula* 3,910 paralog pairs were detected [81], with 2,140 of these (4,185 unique genes) passing our gene expression and variance filtering criteria. These paralogs were significantly under-represented for eGenes, with 1,078 of the 4,185 genes having at least one associated eSNP (hypergeometric test, p = 0.0004). Comparing the expression correlation of paralog pairs to that of random gene pairs showed that paralogs exhibited conserved regulation (permutation test, p < 0.001). We compared the expression correlation distributions of paralog pairs containing 0, 1, and 2 eGenes (Fig 4) and found that a higher number of eGenes in a pair was associated with lower expression correlation (linear model 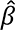 = -**0.06**, p < 2.2x10^-16^). Excluding paralogs did not alter the fact that eGenes had significantly lower network connectivity than non-eGenes.

**Fig 4.**
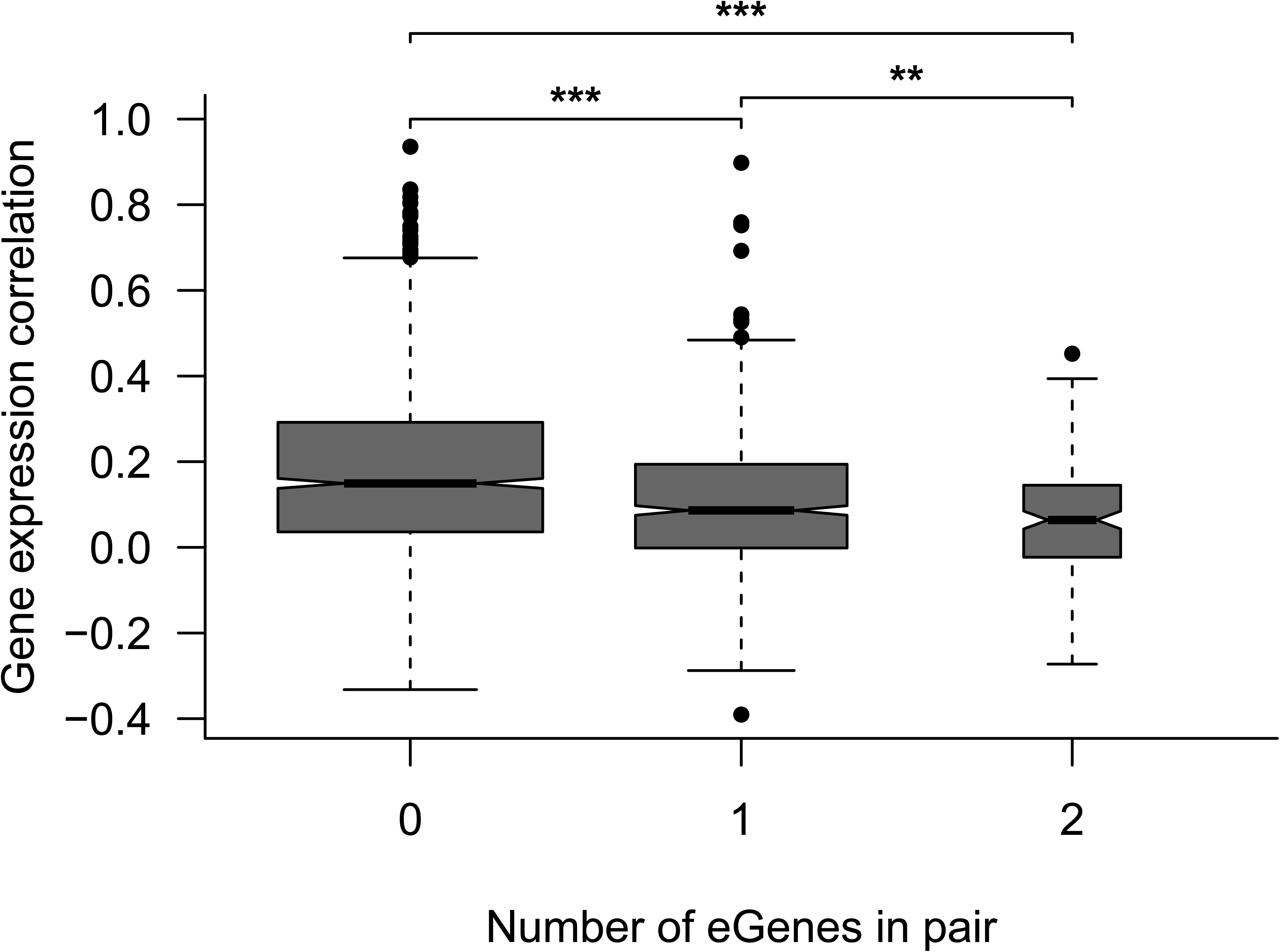
Correlation within paralog pairs as a function of the number of eGenes in the paralog pair. The widths of the boxes are proportional to the number of genes in each set. The mean correlations for paralog pairs with 0, 1, or 2 eGenes were 0.17, 0.10, and 0.06, respectively. The pairwise significance of a Mann-Whitney test is indicated by asterisks: ***p < 0.001, **p < 0.01.

### The rate of sequence evolution is associated with network topology

To assess whether genes with and without eQTLs (eGenes vs. non-eGenes) are experiencing different levels of selective constraint, we assessed nucleotide diversity (θ_π_) and the site frequency spectrum of segregating mutations using Tajima’s D in different genomic contexts. We found that, irrespective of whether eQTLs were local or distant, eGenes exhibited significantly higher genetic variation than non-eGenes (Fig 5A, S1 Table). Furthermore, although Tajima’s D values are overall negative in *P. tremula*, likely reflecting a historical range expansion [63], eGenes exhibited significantly higher Tajima’s D values compared to non-eGenes (Fig 5C, S1 Table), again regardless of whether the associated eQTL was local or distant. As such, non-eGenes appear to be experiencing stronger selective constraint than eGenes. These patterns were consistent for the different genetic contexts considered (S16 Fig), likely reflecting the effects of linked selection [63]. To test this hypothesis we looked for differences in selection efficacy on protein sequence evolution between eGenes and non-eGenes by examining the ratios of intraspecific nonsynonymous to synonymous polymorphisms (θ_0-_f_old_/θ_4-_f_old_) and interspecific nonsynonymous to synonymous substitutions (d_N_/d_S_) across these genes (using *P. trichocarpa* as an outgroup). Both θ_0-fold_/θ_4-fold_ and d_N_/d_S_ estimates were significantly lower in non-eGenes compared to eGenes (Fig 5E,G, S1 Table), likely reflecting stronger purifying selection acting on non-eGenes. As has been reported previously for *Capsella grandiflora* [27] and a wild population of baboons [83], we observed a negative correlation between minor allele frequency and eQTL effect size (S17 Fig). We examined this relationship in permuted data, which revealed an excess of low MAF SNPs in the permuted data compared to the original data (S18 Fig). This is in contrast to the results in *Capsella* [27], where the reverse was observed. However, in both [27] and in our results there was a consistent negative relationship between MAF and effect size. The comparative differences in MAF distributions for real and permuted data between the two studies may largely result from statistical differences in how the test statistic and p-values were calculated, but may also reflect differences in the population genetics of the two systems. One concern is that this relationship may result from a higher false positive rate at lower MAF due to the concomitant decrease in sample size. To address this concern, we performed a subsampling analysis, similarly to [27], to remove the effect of MAF and examined the correlation of effect sizes estimated in the original and sub-sampled datasets. The high correlation observed (S19 Fig) suggests that the observed negative relationship between effect size and MAF is not artefactual.

**Fig 5.**
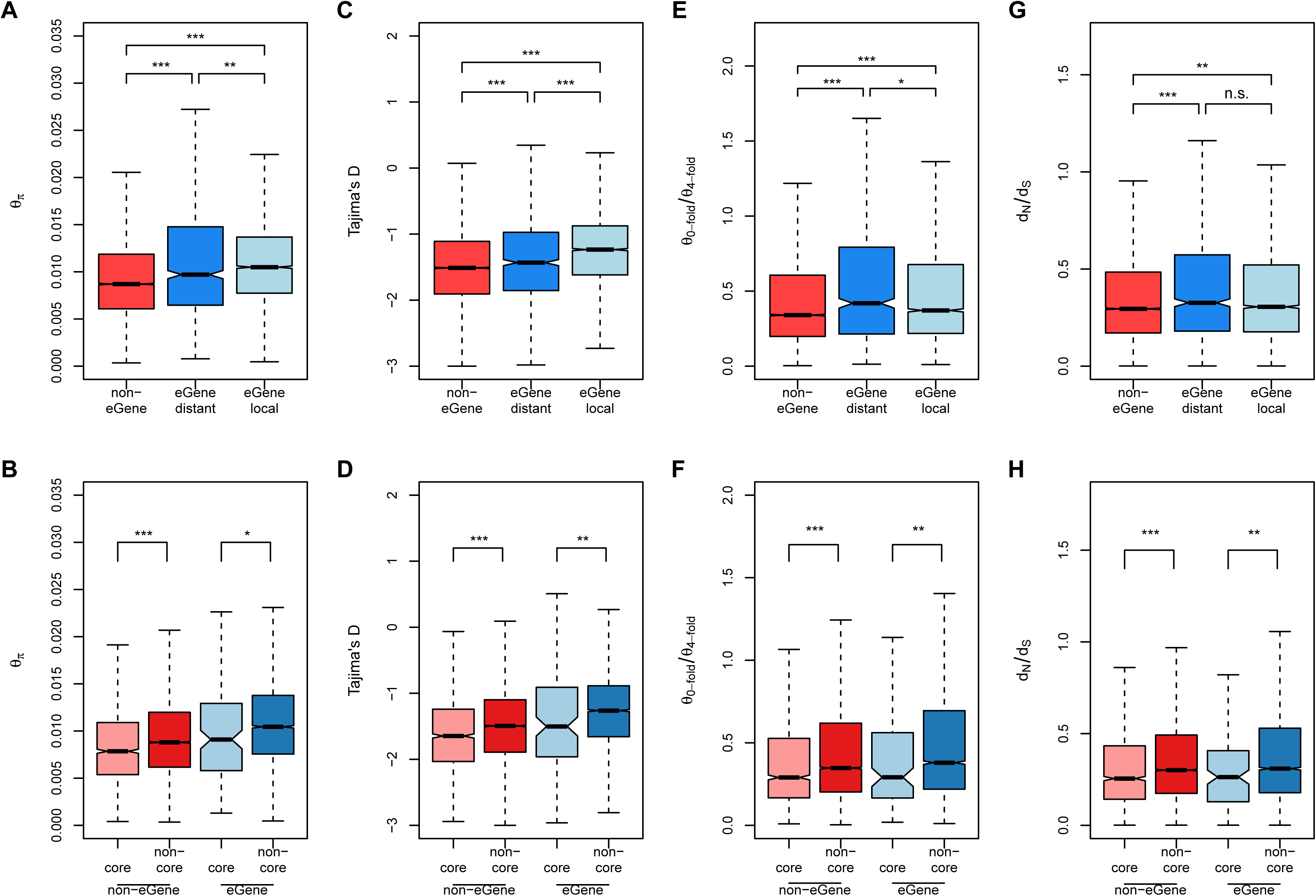
Measures of sequence diversity and divergence. Nucleotide diversity (A,B), Tajima’s D (C, D), θ_0-fold_/θ_4-fold_ (E,F) and d_N_/d_S_ (G,H) are compared between eGenes (with local eQTLs or with only distant eQTLs) and non-eGenes, as well as core and non-core genes from the gene expression network. Significance between each pair of gene categories was evaluated using Mann-Whitney tests and significance is indicated by asterisks above the boxplot: ***p < 0.001, **p < 0.01, *p < 0.05, n.s. p >0.05.

To further examine the relationship between network topology and sequence evolution, we contrasted genes located in module cores and those not (core vs. non-core). Independent of whether a gene was an eGene or not, core genes had significantly lower levels of genetic diversity and Tajima’s D values compared to non-core genes (Fig 5B,D). In addition, core-genes also had significantly reduced ratios of non-synonymous to synonymous polymorphisms (θ_0-fold_/θ_4-fold_) and substitutions (d_N_/d_S_) (Fig 5F,H). Again, these patterns were consistent across different genomic contexts (S16 Fig). Taken together, these results suggest that genes in network module cores experience reduced rates of molecular evolution due to stronger purifying selection, i.e. selective removal of deleterious mutations, and are therefore evolving under stronger selective constraint compared to non-core genes. Stronger purifying selection on mutations within core genes is likely driven by stronger stabilising selection of gene expression noise or modulation acting to maintain the optimal level of expression in core, compared to peripheral, genes [84].

As the sequence evolution of a given gene is known to correlate with different factors [85], such as gene expression level or variance, the evolutionary age of a gene [86,87] and, as we show, the presence or absence of an eQTL and the topology of the co-expression network (S20 Fig), we performed analyses to ascertain their relative roles in determining patterns of sequence evolution. Owing to the collinearity of various characteristics of expression (S20 Fig), we performed principal component analysis (PCA) on representative gene expression measures (before hidden confounder removal) to examine the extent to which these measures were interdependent. This analysis revealed that PC1, which explained 37.03% of the variation in these five measures, was mainly dominated by the connectivity of genes in the co-expression network and whether they are located within network module cores or not (Fig 6A). Gene expression variance and eGene status showed a strong influence on PC2 (Fig 6A) while expression level contributed largely to PC3 (Fig 6A).

**Fig 6.**
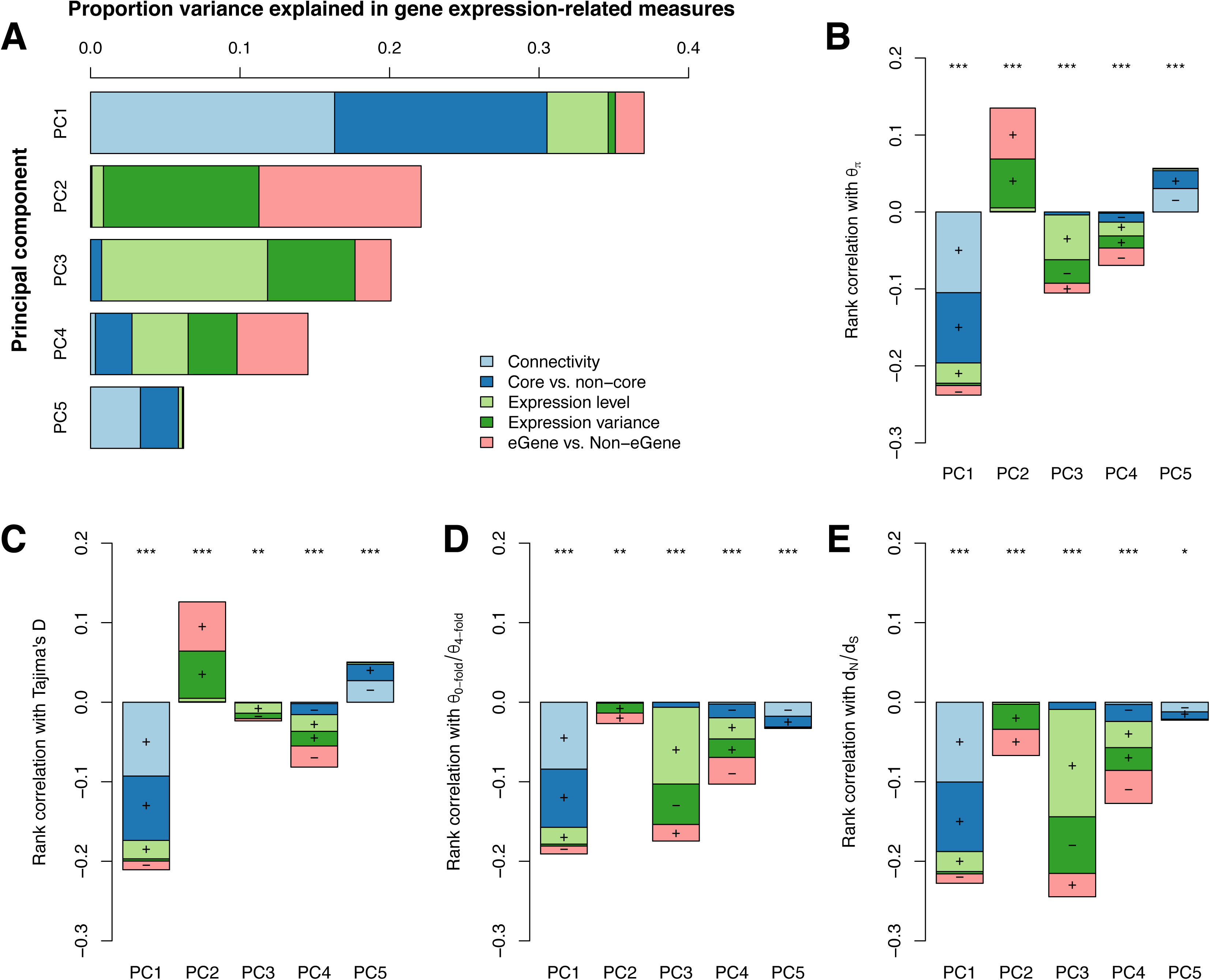
Associations between metrics of gene expression and sequence evolution in *Populus tremula*. (A) Percentage variance explained by five principal components comprising co-expression network connectivity, genes in network cores or not (Core vs. non-core genes), gene expression levels, gene expression variance and genes with eQTLs or not (eGenes vs. non-eGenes). Colour shadings depict the proportion contribution of each gene expression measure to each principal component. (B-E) Spearman’s rank correlations between PCs and four metrics of sequence evolution: nucleotide diversity (θ_π_), Tajima’s D, 0_0-fold_/0_4-fold_ and d_N_/d_S_. Stacking of the barplots shows the relative contribution of each gene expression measure to each PC. Plus or minus indicates the direction of correlation for individual variable on the corresponding PCs. Asterisks indicate significant correlations, *\*\*\*P* < 1e-5, *\*\*P* < 0.001,* P<0.05, ns=not significant (*P* >0.05).

In order to test whether the correlations between sequence evolution and various characteristics of expression are independent of one another, we calculated correlations (Spearman’s rank) between sequence evolution rates and PCs. The rates of sequence evolution over both short (represented by within-species diversity: θ_π_, Tajima’s D and θ_0-fold_/θ_4-fold_) and long (represented by between-species divergence, d_N_/d_S_) timescales showed significantly negative correlations with the connectivity and core status of genes in the co-expression network (PC1) (Fig 6B-E). This indicates that the connectivity of genes within the network, both globally and within the local context of expression modules (core status), are key factors associated with the rates of sequence evolution. PC2, which largely reflected gene expression variance showed significant positive correlations with patterns of genetic diversity within species (θ_π_ and Tajima’s D), and had significant, although relatively weak, negative correlation with the rate of protein sequence evolution (Fig 6B-E). In addition, in accordance with other studies [88], gene expression level (largely represented by PC3) showed significantly negative correlations with the rate of protein sequence evolution (Fig 6B-E). We then performed partial Spearman correlation, aiming to estimate the relationship between two variables while controlling for other variables, between sequence evolution measures and the five measures of gene expression used in the above PCA (Table 1). We found the correlation between sequence evolution and connectivity of genes within the co-expression network persists, although slightly lower, after accounting for the effects of gene expression level and other factors (Table 1).

**Table 1.** Summary of the correlation coefficients (Spearman’s rank correlation coefficient) between measures of sequence evolution and gene expression-related measures.

| | $\theta_{\pi}$ | | Tajima's D | | $\theta_{0\text{-fold}}/\theta_{4\text{-fold}}$ | | $d_N/d_S$ | |
| --- | --- | --- | --- | --- | --- | --- | --- | --- |
|  | Pairwise | Partial<sup>a</sup> | Pairwise | Partial<sup>a</sup> | Pairwise | Partial<sup>a</sup> | Pairwise | Partial<sup>a</sup> |
| Connectivity | -0.2197*** | -0.1440*** | -0.1929*** | -0.1164*** | -0.1352*** | -0.0417*** | -0.1529*** | -0.0520*** |
| Core vs. Non-core | -0.0848*** | -0.0045ns | -0.0938*** | -0.0153* | -0.0652*** | -0.0146* | -0.0588*** | 0.0009ns |
| Expression level | -0.1312*** | -0.0729*** | -0.0817*** | -0.0370*** | -0.2213*** | -0.1914*** | -0.2792*** | -0.2441*** |
| Expression variance | 0.1238*** | 0.1263*** | 0.0580*** | 0.0587*** | 0.0356*** | 0.0320** | 0.0302** | 0.0241** |
| eGene vs. Non-eGene | 0.1677*** | 0.1159*** | 0.1869*** | 0.1419*** | 0.0540*** | 0.0457*** | 0.0309** | 0.0265** |

<sup>a</sup> Partial correlations between two specific variables after controlling for all other gene-expression related measures.
ns=not significant (P>0.05)
\*P<0.05
\*\*P<0.001
\*\*\*P<1e-5

## Discussion

Our primary aim was to determine the evolutionary forces maintaining genetic variance associated with gene expression variation and to determine their relationship to the associated gene co-expression network topology within a wild-collected, outbreeding population of the forest tree *Populus tremula*. We first established that, in common to results in other species [4–16,52], there is prevalent heritability in gene expression levels, with 17% of the genes (5,924) having *H*^2^ > 0.5. To examine the genetic architecture of heritable gene expression variation within the population, we performed eQTL mapping, from which we identified more local than distant eQTLs, with local eQTLs explaining significantly more of the variance in gene expression than distant eQTLs (Fig 2A), which is similarly in agreement with previous studies across a range of species [34,47–50]. Although each eSNP was typically associated with only a single gene, many genes were associated with more than one unique eSNP, indicating that numerous loci influence the expression of genes with an associated eQTL. For a large proportion of eGenes, the identified set of eSNPs explained a relatively high proportion of the heritable expression variation (median 1.0 and s.d. 2.3; S21 Fig). While it appeared that a single eSNP often accounted for a large proportion of the explained heritable variance, due to linkage it is not possible to determine the relative contribution of each eSNP within a haplotype block, with the apparent contribution of individual eSNPs largely resulting from the order in which they are entered into the statistical model. While eGenes had higher *H*^2^ than non-eGenes, as is expected, there were, nonetheless, 2,780 non-eGenes with *H^2^* >0.5, potentially reflecting that the expression variance of these genes is controlled by many eSNPs of small effect size that we lacked the power to detect.

To gain insight into how eQTLs may be influencing expression of the associated eGene, we examined their genomic context. Local eQTLs were most frequently located in regulatory regions (Fig 2D), with UTRs having the highest density of local eSNPs (∼1.5 eSNP per kbp) followed by flanking regions (2 kbp up- and downstream, ∼1 per kbp) and introns (∼0.75 per kbp), which is consistent with previous findings in natural populations of humans [76,78], Drosophila [79] and Capsella grandiflora [27], likely indicates that these loci cause regulatory changes in the transcriptional dynamics of the gene.

To address our primary question, we used the gene expression values to construct a co-expression network. Compared to networks more typical of systems biology studies, for example the *P. tremula* exAtlas network [64], where samples originated from different tissues of a single genotype, the pairwise expression correlations underlying our co-expression network were low. Despite this, the network displayed typical characteristics, being scale-free with hubs and distinct modules [80]. To determine whether network topology is related to the evolutionary history of its component genes, we examined correlations between network connectivity and rates of sequence evolution. Scale-free gene co-expression networks are defined as robust, or buffered, to the effects of random mutations. As there are few highly connected genes (which are important determinants of the observed co-expression structure), a random mutation would be unlikely to affect such a gene. In comparison, as the majority of genes have low connectivity, a random mutation is more likely to affect such a low connectivity gene, which have few connections to other genes in the network and the effect of this mutation will therefore be minimal (i.e. the overall network is buffered from, or robust to, the mutational effect) [89]. In network modules, where many genes share the same expression pattern, a single eSNP modulating the expression of a central regulator would be sufficient to induce similar expression variation (i.e. co-expression) in the set of connected genes within that network module. The buffered characteristic of the network would therefore hold true even if all genes within the network are exposed to the same evolutionary history – i.e. that all are equally likely to accumulate mutations. However, natural selection may interact in combination with network topology, for example to prevent the accumulation of mutations within specific genes [90]. If the distribution of mutations is not random across the network, such an interaction can be inferred. In our data, genes in module cores (the set of the most highly connected genes within a module) were under-represented for eGenes and, respectively, those in the network periphery were enriched for eGenes (Fig 3B), suggesting that polymorphisms associated with natural variation in expression are more likely to affect genes of low connectivity. More generally, eGene connectivity was negatively correlated with variance explained by the associated eSNPs (S15 Fig). It has similarly been reported that hubs in human protein-protein interaction networks are less likely to be associated with a detectable eQTL and that the effect size of eQTLs is negatively correlated with connectivity in the protein-protein interaction network [91]. Furthermore, connectivity and core status contributed primarily to PCs that were independent to expression level (Fig 6A), with connectivity being correlated negatively with the rates of sequence evolution at both short (Fig 6B-C) and long (Fig 6D-E) timescales. In comparison, expression level exhibited low correlation with sequence evolution at short timescales, but higher negative correlations over long timescales (Fig 6B-E). Genes with high co-expression connectivity, mostly those within network module cores, have significantly reduced genetic diversity and exhibit reduced rates of protein evolution compared to more peripheral genes (Fig 5). This suggests that core genes, which represent genes of high potential effect, have evolved under stronger evolutionary constraint than genes in the periphery of the network. As genes in module cores generally have higher levels of pleiotropy and lower levels of dispensability [58], they are consequently more constrained against changes in both gene expression and protein sequence compared to genes in the periphery of the network [92]. Furthermore, we show that eGenes are overrepresented among non-core genes of the network and that they have experienced lower levels of purifying selection relative to non-eGenes, corroborating a relaxation of both expression and coding constraints for these genes. Relaxed selection of peripheral eGenes is thus expected to result in an accumulation of weakly deleterious mutations that will segregate as intra-specific polymorphisms. Consistent with these observations, we found that conserved genes (i.e. old genes) were significantly over-represented in network cores while P. tremula specific genes (i.e. young genes) were under-represented. This is in agreement with previous studies that have shown that evolutionarily ancient genes tend to be more central in regulatory networks, with increased constraints on expression and fewer associated eQTLs [87,86]. Taken together, these results indicate how the co-expression network can be buffered against large perturbations via constraint of core genes while enabling flexibility and adaptation by tolerating an accumulation of mutations within the network periphery.

The Salicaceae lineage underwent a relatively recent (58 million years ago) whole-genome duplication shared by all member species and that remains represented by a large number of paralogous gene pairs in the genomes of *Populus* species [82]. If many of these duplicated genes are functionally redundant or in the process of diverging, one would expect them to be overrepresented for eGenes as sub- or neo-functionalisation requires derived SNPs to drive expression or coding divergence. However, we saw an under-representation of eGenes in paralog pairs, suggesting that the SNPs that initially induced expression divergence have reached fixation. Interestingly, we found progressively lower expression correlation between paralog pairs containing zero, one or two eGenes (Fig 4), indicating that a subset of paralogs are still undergoing sub- or neo-functionalisation, with their associated eSNPs driving expression divergence.

## Conclusion

We identified substantial heritability of gene expression within a natural, out-breeding population, of *P. tremula*. Polymorphisms associated with expression variance were most frequently located within regulatory regions, suggesting that they act by inducing expression variance. The gene co-expression network displayed typical characteristics, being scale-free i.e. containing a small number of highly connected genes. The most highly connected genes within module cores were underrepresented by eQTLs, with a negative correlation between eQTL effect size and network connectivity. In contrast, the network periphery was enriched for eQTLs, suggesting higher selective constraint on expression variance within the network core (stabilising selection) and relaxed constraint within the periphery. Integration of the eQTL and population genetics analyses with characteristics of the associated gene co-expression network highlight that the context of a gene within the co-expression network appears to be an important determinant of the evolutionary dynamics of transcribed loci. Our results point towards stronger selection acting on network core genes compared to genes in the periphery of the co-expression network, with a negative correlation between rates of sequence evolution and gene connectivity. Taken together, this suggests that highly connected genes within the core of the co-expression network derived from flushing buds of P. tremula have experienced stronger purifying selection than those in the network periphery, the action of which is associated with higher stabilising selection of gene expression variance for highly connected genes.

## Materials and Methods

### Samples

We collected branches form the SwAsp collection common garden in north of Sweden on 27th May 2012, before natural bud break but as close to the point of natural spring bud break as possible. Branches were placed in the greenhouse facility at the Umeå Plant Science Centre under conditions selected to induce rapid bud break (24 h light, temperature of 20 °C and humidity 50-70 %). At a defined point of emergence (S22 Fig) buds were harvested, flash frozen in liquid nitrogen and stored at -80 ° C until used for RNA isolation. Only terminal buds were sampled (i.e. no lateral buds were included). The time from the day branches were placed in the greenhouse until bud flush sampling ranged from one day to eight days (S23A Fig) and there was a high, positive correlation to bud flush date recorded in the field for the same year (S23B Fig; r=0.776, p < 2.2×10^-16^). As has previously been reported [93], there was very low *Q*_ST_ for bud flush, either in the field or for the greenhouse material ^(^*Q*_ST_ 0.13 and 0.07 respectively), however *H^2^* was high (*H^2^* = 0.82 and 0.71 respectively).

### RNA isolation

One to two buds per clonal replicate were ground using one 3 mm stainless steel bead (Qiagen, Redwood city, USA) in Corning^®^ 96 well PP 1.2 ml cluster tubes (Sigma-Aldrich, St. Louis, USA) using a Mixer Mill MM400 (Retsch, Haan, Germany) at 20 Hz for 2 x 15 sec. Total RNA was extracted from all samples according to [94] with the omission of the L spermidine. Buffer volumes were adjusted according to starting material (70 - 130 mg). RNA isolation was performed using one extraction with CTAB buffer followed by one chloroform: isoamyl alcohol IAA (24:1) extraction. All other steps were performed as in [94]. DNA contamination was removed using DNA-free™ DNA removal Kit (Life Technologies, Carlsbad, USA). RNA purity was measured using a NanoDrop 2000 (Thermo Scientific, Wilmington, USA) and RNA integrity was assessed using the Plant RNA Nano Kit for the Bioanalyzer (Agilent Technologies, Santa Clara, USA).

### RNA-Sequencing and analysis

RNA-Sequencing was performed as in [74]. Briefly, paired-end (2 × 100 bp) RNA-Seq data were generated using standard Illumina protocols and kits (TruSeq SBS KIT-HS v3, FC-401-3001; TruSeq PE Cluster Kit v3, PE-401-3001) and all sequencing was performed using the Illumina HiSeq 2000 platform at the Science for Life Laboratory, Stockholm, Sweden. Raw data is available at the European Nucleotide Archive (ENA, [95]) with accession number ERP014886, and the normalized and filtered gene expression data matrix is available at the PlantGenIE FTP resource (ftp://plantgenie.org/Publications/Maehler2016/).

RNA-Seq FASTQ-files were pre-processed and aligned to v1.0 of the *P. tremula* reference genome [81] as in [96]. In short, reads were quality and adapter trimmed using Trimmomatic v0.32 [97], rRNA matching reads were filtered using SortMeRNA v1.9 [98], reads were aligned to the v1.0 *P*. *tremula* reference genome using STAR 2.4.0f1 [99] and read counts were obtained using htseq-count from HTSeq [100]. FastQC [101] was used to track read quality throughout the process. Normalised gene expression values were obtained by applying a variance stabilising transformation (VST) to the raw counts from HTSeq, as implemented in the DESeq2 R package [102].

### Gene expression *H*^2^ and *Q*_ST_

We calculated repeatability as an assumed upper bound estimate of broad sense heritability of gene expression (see [103] for discussion) from the variance estimates in our data according to the equation

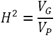

where *V*_G_ is the genetic component of the variance calculated as the expression variance between genotypes for a particular gene (*i.e*. variance among individual means) and *V*_P_ is the total phenotypic variance calculated as the sum of *V*_G_ and *V*_E_, where *V*_E_ is the environment component of the variance calculated as the expression variance within genotypes for a particular gene (i.e. the mean variance among clonal replicates). Point estimates of *H*^2^ were obtained using the repeatability function from the heritability v1.1 R package [104]. The significance of the broad sense heritability was estimated by using an empirical null model where heritabilities were based on random genotype assignments. For each gene, 1,000 permutations were performed and empirical p-values were calculated using the empPvals function in the qvalue 2.6.0 R package.

Population differentiation *Q*_ST_, [105]) was calculated as

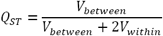

where *V*_between_ is the variance among populations and *V*_within_ is the residual genetic variance among genotypes within populations as computed using the lmer function from the lme4 v1.1.12 R package [106] using the formula

**expression ∼ 1 + (1 | population) + (l|clone)**

where *expression* is the expression of a gene, *population* is a factor representing the population of each sample, and *clone* is a factor representing genotype replicates. As we use repeatability as an upper bound estimate of *H*^2^, our *Q*_ST_ estimates are conservative [107]. Similar to the broad sense heritability, the significance of the *Q*_ST_ was estimated using permutation testing where genotype labels were shuffled among the populations 1,000 times. Due to the characteristics of the data and the method, many of the permutations resulted in undefined QST estimates (an average of 511 missing values per gene). Thus, the empirical p-values for the genes were based on different number of null values.

### Hidden confounder removal

Genotype mean gene expression data was adjusted for hidden confounders before mapping eQTLs and constructing the co-expression network. Hidden confounders in the gene expression data was accounted for by regressing out the 9 first principal components (PCs) of the gene expression data [75–77]. The number of components to remove was determined by running the eQTL mapping with 0 to 20 PCs removed and selecting the number of components that yielded the largest number of significant eQTLs (Benjamini-Hochberg p < 0.05) (S5 Fig). This approach was based on the assumption that the number of identified eQTLs would increase if the removed PCs were removing unwanted, systematic variation (i.e. noise) rather than informative biological variation [75–77,108,109].

### eQTL mapping

eQTL mapping was performed by associating genotype mean gene expression levels with biallelic SNPs using the R package Matrix eQTL v2.1.1 [110]. The corresponding raw resequencing data for all SwAsp genotypes is available at the NCBI SRA resource as BioProject PRJNA297202 (SRA: SRP065057) and the resultant Variant Call Format (VCF) file used for association mapping is available from [65]. Before performing association mapping, genes were filtered on variance so that only genes with a gene expression variance above 0.05 were included. SNPs were also filtered on minor allele frequency (MAF) and major genotype frequency (MGF); any SNPs with MAF < 0.1 or MGF > 0.9 were excluded to avoid spurious associations. A motivating example for the MGF filtering can be seen in S24 Fig. We also generated an LD-trimmed SNP set by removing one SNP from each pair of SNPs with a between SNP correlation coefficient (*r*^2^) >0.2 in blocks of 50 SNPs using PLINK v1.9 [111], yielding 217,489 independent SNPs that were retained for analyses of population structure. The first genotype principal component based on the set of independent SNPs was used as a covariate in the linear model used by Matrix eQTL to account for the weak signature of population structure. Permutation testing was used to determine eQTL significance whereby genotype sample labels were permuted 1,000 times and the maximum absolute t-statistic from Matrix eQTL was recorded for each expressed gene across all SNPs for each permutation, resulting in 1,000 random t-statistics being collected for each gene. Empirical p-values were calculated for each eQTL in the observed data using the permuted t-statistics for the observed eGene with the empPvals function in the qvalue v2.4.2 R package [112], and q-values (empirical FDR) were calculated with the qvalue function in the same package.

When determining the genomic context of eSNPs, there were some cases where introns overlapped exons as a result of overlapping gene model being present on the same strand. These 41 eSNPs were discarded from the counting. Another type of overlap that was discarded were cases where an eSNP overlapped a gene feature, but no sub-feature inside that gene (e.g. UTR, exon or intron). These 1961 eSNPs were excluded from the counting. Since many of the features overlap (e.g. exon and untranslated regions), the priority for counting was untranslated region, exon/intron, upstream/downstream and intergenic.

Permutation tests involving eGenes and non-eGenes were performed by shuffling eGene assignments among the 22,306 genes that were considered for eQTL mapping. This was repeated 10,000 times and empirical p-values were calculated using the empPvals function in the qvalue v2.6.0 R package. The minor allele frequency (MAF) was compared between eSNPs (both local and distant) and all SNPs throughout the genome. To test for the signature of selection on eQTLs, we also estimated the correlation between minor allele frequency and eQTL effect size.

Permutation analysis of the minor allele frequency distribution was performed by generating 20 sets of 150,000 random SNPs from the set of MAF and MGF-filtered SNPs (sampled with replacement). eQTL mapping was performed for each of these sets, and in addition, the sample labels of the genotype data was shuffled 50 times for each SNP-set and consequently used for eQTL mapping, resulting in a total of 1,000 permuted eQTL sets.

For the subsampling analysis, the significant eSNPs from the original eQTL mapping were subsampled in order to fix the minor allele frequency. It was decided to fix the minor allele frequency at 25% and use 40 samples as this retained 136,942 of the original 147,419 eSNPs (93%). The actual subsampling was done by dropping samples recursively until the minor allele frequency fell within ±2 percentage points of the desired frequency. This was done using our custom program vcfsubsample (https://github.com/maehler/vcfsubsample).

### Co-expression

The R package WGCNA v1.51 [113] was used for constructing a co-expression network. The input gene expression values were per-gene genotype means corrected for hidden confounders (the same data used for eQTL mapping). We chose to use the unsigned network type for this study with the motivation that we did not want to discard negative relationships. By looking at this from an eQTL perspective, an eSNP can be positively associated with one gene while negatively associated with another. The relationship between these two genes would be missed if we used the signed or the signed-hybrid approaches. Using the unsigned approach, we assure that genes with strong negative correlation end up in the same network modules. A soft thresholding power of 5 was used to calculate adjacencies. The topological overlap matrix (TOM) was generated using the TOMsimilarity function with the signed approach in order to take negative edges into account (see [113] for details). In order to identify network modules, hierarchical clustering was applied to the TOM dissimilarity matrix (1 – TOM) and the resulting dendrogram was divided into modules using the cutreedynamic function. The connectivity of the network was then defined as the adjacency sum for each node, i.e. the weights of the edges that are connected to this node. This concept was applied to modules as well to obtain measures of intra- and inter-modular connectivity, i.e. the connectivity based on edges connecting the gene with other genes inside the same module, and connectivity based on edges connecting the gene with genes outside of the module.

To define the periphery of the network we applied a hard edge-threshold to the network where only gene-pairs with an absolute Pearson correlation > 0.4 were linked, which corresponded to the top 0.1% most correlated gene pairs. Genes were then classed as peripheral if they linked to only one other gene.

Conserved genes and *P*. *tremula*-specific genes were inferred from gene family analysis based on protein sequence similarity available on the PopGenIE FTP server [114]. In short, two rounds of TribeMCL were run using BLASTP with an E-value threshold of 10^-5^ where the first round was all-vs-all while the second was all-vs-all inside each gene family.

Transcription factors were identified by BLASTing the the *P. tremula* protein sequences against the protein sequences of transcription factors in *P. trichocarpa* with an E-value threshold of 10^-10^ and taking the best hit for each *P. trichocarapa* transcription factor. The *P. trichocarpa* transcription factor annotations were taken from Plant Transcription Factor Database v3.0 [115].

All permutation tests based on the network were performed by computing the measure of interest on the original network, and then performing the same computation on 10,000 networks were the gene labels were shuffled. Empirical p-values were calculated using the empPvals function in the qvalue v2.6.0 R package [112].

Functional enrichment was tested using the R package topGO 2.24.0. As background the 22,306 genes included in the network construction were used.

### Population genetic analysis

To estimate nucleotide diversity (θ_π_) and Tajima’s D [116] among genes, we only used the reads with mapping quality above 30 and the bases with quality score higher than 20 for the estimation of θ_π_ and Tajima’s D by ANGSD [117]. Genes with less than 50 covered sites left from previous quality filtering steps (Wang et al. In prep) were excluded. In addition, θ_π_ and Tajima’s D were also calculated for seven site categories of each gene: 1kbp upstream of genes, 1kbp downstream of genes, 3’ UTR, 5’ UTR, intron, 0-fold non-synonymous and 4-fold synonymous sites within genes. We further compared the estimates of θ_0-fold_/θ_4-fold_ between different categories of genes. We used the transcript with the highest content of protein-coding sites to categorize the seven genomic features within each gene. Finally, we estimated the ratio of non-synonymous substitution and synonymous substitution rate (d_N_/d_S_) using gKaKs pipeline [118] with codeml method [119] for a total of 33,039 orthologous genes that were determined by best BLASTp sequence homology searches between *P. tremula* and *P. trichocarpa* [81,114]. Significance for the above statistical measurements between each pair of gene categories was evaluated using Mann-Whitney tests.

To test for the main effects of various gene expression factors (including connectivity within co-expression network, core vs. noncore, presence or absence of an eQTL, gene expression level and variance) on the rate of sequence evolution, we first performed principal component analysis (PCA) of the five gene expression-related variables to examine the extent to which they were interdependent. To further handle the problem of collinearity between them, the PCs were calculated. We then examined the Spearman’s rank correlation coefficient of each PC with the parameters of sequence evolution. The data were scaled before doing PCA, and the computation of PCs was implemented in the “prcomp” function of the statistical software package R 3.2.0.

Finally, to account for the autocorrelation between gene expression measurements, we estimated partial Spearman correlations between sequence evolution parameters and each of gene expression related variables while controlling for other variables. The partial correlation were performed with R package ppcor.

### Analysis script availability

Relevant processed data and R transcripts are available on the PlantGenIE FTP resource (ftp://plantgenie.org/Publications/Maehler2016/).

## Acknowledgements

We thank Dr Kathryn Robinson for her help sampling material, for providing environmental data and for informative discussion. The authors also would like to acknowledge support from the Science for Life Laboratory, the National Genomics Infrastructure (NGI), and the Uppsala Multidisciplinary Center for Advanced Computational Science (UPPMAX) for providing assistance in massive parallel sequencing and computational infrastructure. The authors wish to thank the three reviewers, who offered invaluable feedback and suggestions during the review process. Their input substantially improved the presentation and interpretation of the study and we are grateful for the time they dedicated to this.

## Supporting information

**S1 File. GO enrichment analysis results**. Contains GO enrichment results for local and distant eGenes, network module cores, and subsets of genes with extreme *H*^2^ and *Q*_ST_ values.

**S2 File. Various gene statistics.** Contains statistics for all genes, as well as all genes included in the generation of the co-expression network and the eQTL mapping.

**S1 Table. Statistic summary (median and central 95% range) for four measures of sequence evolution:** θ_π_, Tajima’s D, θ_0-fold_/θ_4-fold_ and dN/dS in groups corresponding to Fig 5, and the summary of Wilcoxon rank sum test (location shift and 95% confidence interval) between two groups of estimates shown in Fig 5.

**S1 Fig.**
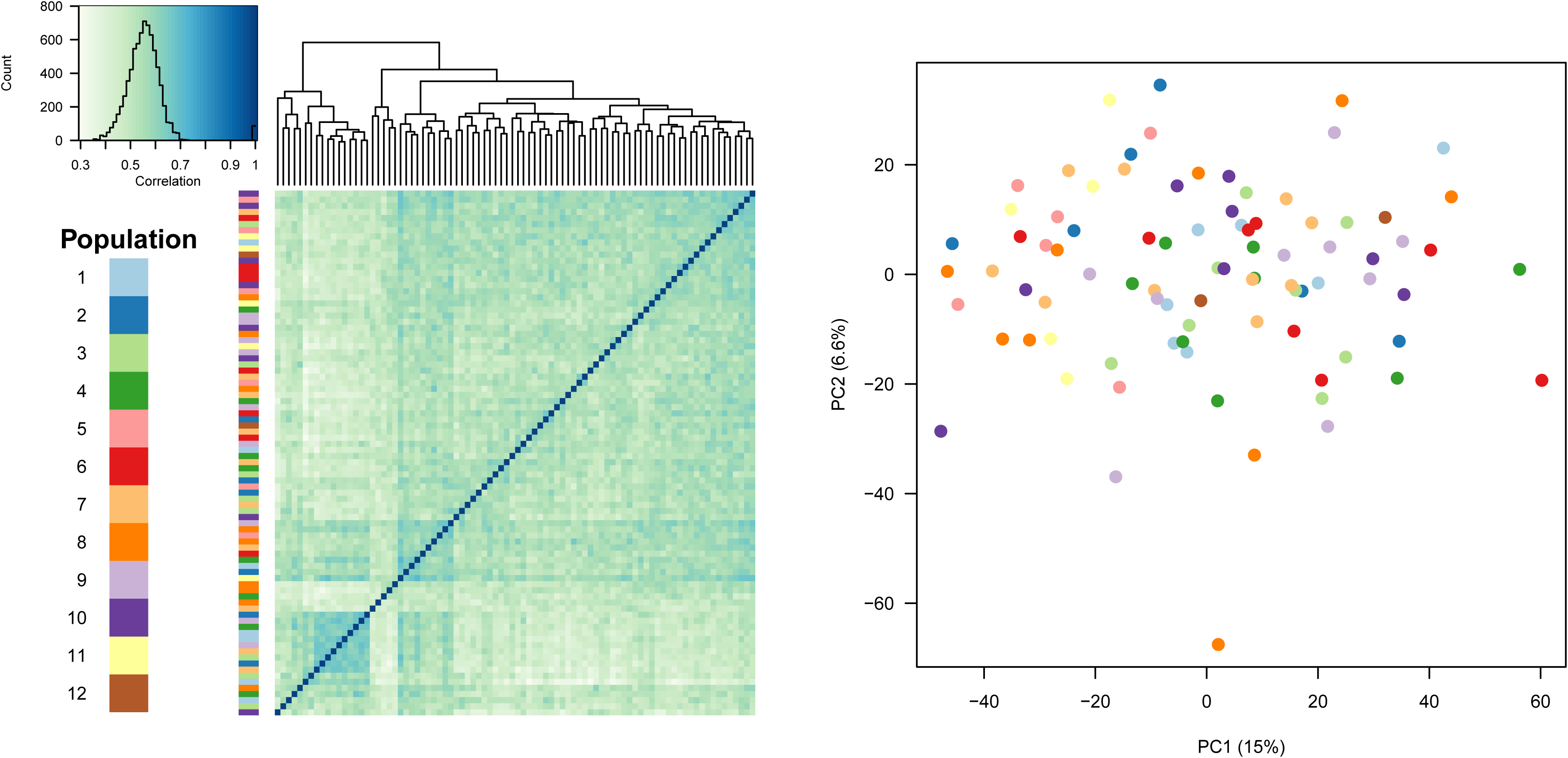
Clustering of genotypes. (Left) Heatmap of the sample correlation matrix based on the 500 most variably expressed genes. Darker colour indicates higher correlation. The coloured bar represents the populations the genotypes belong to. (Right) The two first principal components from a principal component analysis (PCA) based on all genes. Again, colours represent the genotype population. The percentages in the axis labels indicate the amount of variance explained by each component.

**S2 Fig.**
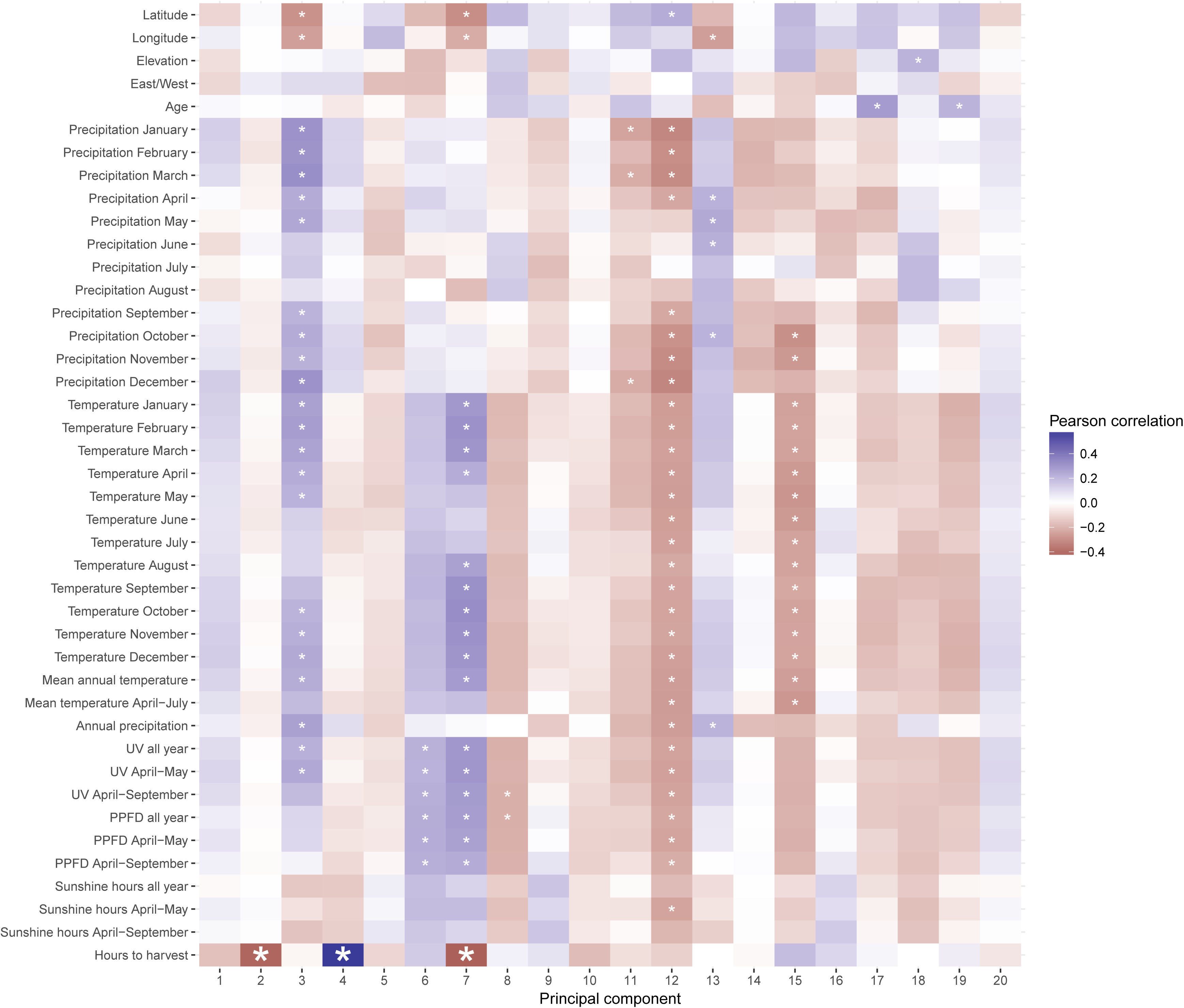
Correlations between gene expression principal components (PCs) and environmental variables. The values in each tile represent the Pearson correlation between the gene expression PC (x-axis) and environmental variable (y-axis). Small asterisks represent a nominal p-value < 0.05 while large asterisks represent Benjamini-Hochberg (BH) adjusted p-values < 0.05. The only factor with significant correlations to expression PCs was “Hours to harvest”, which is the number of hours into the sampling period that the buds were harvested. It was significantly associated with PC4 (BH-adjusted p = 4.6×10^-6^), PC7 (BH-adjusted p = 0.030) and PC2 (BH-adjusted p = 0.033).

**S3 Fig.**
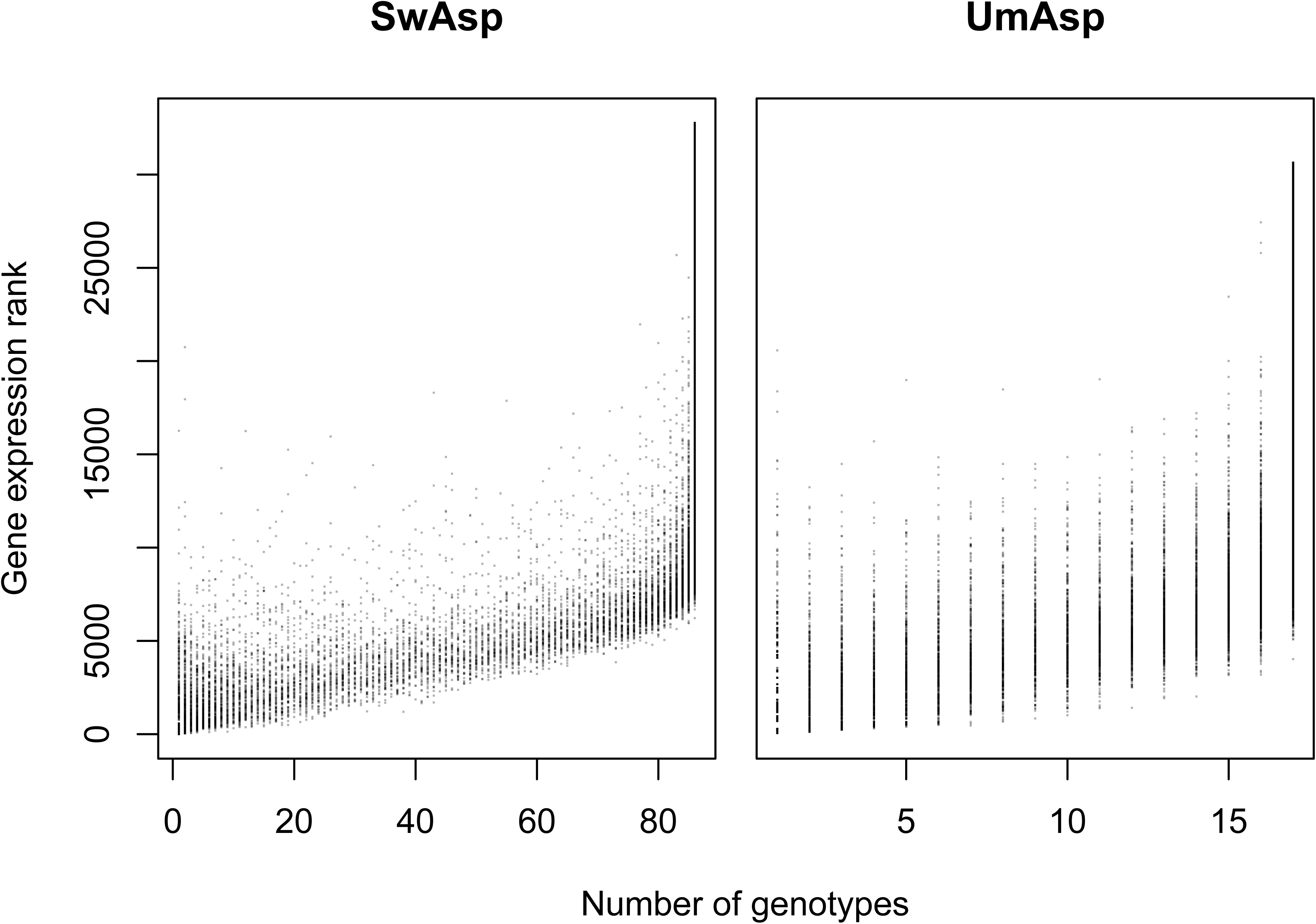
Relationship between gene expression ranks and the number of genotypes with detectable expression of the genes. The number of genotypes that a gene was expressed in was determined by counting the number of genotypes with non-zero expression for each gene. The gene expression ranks were calculated by ranking the mean gene expression values where the mean was calculated only considering samples with non-zero expression.

**S4 Fig.**
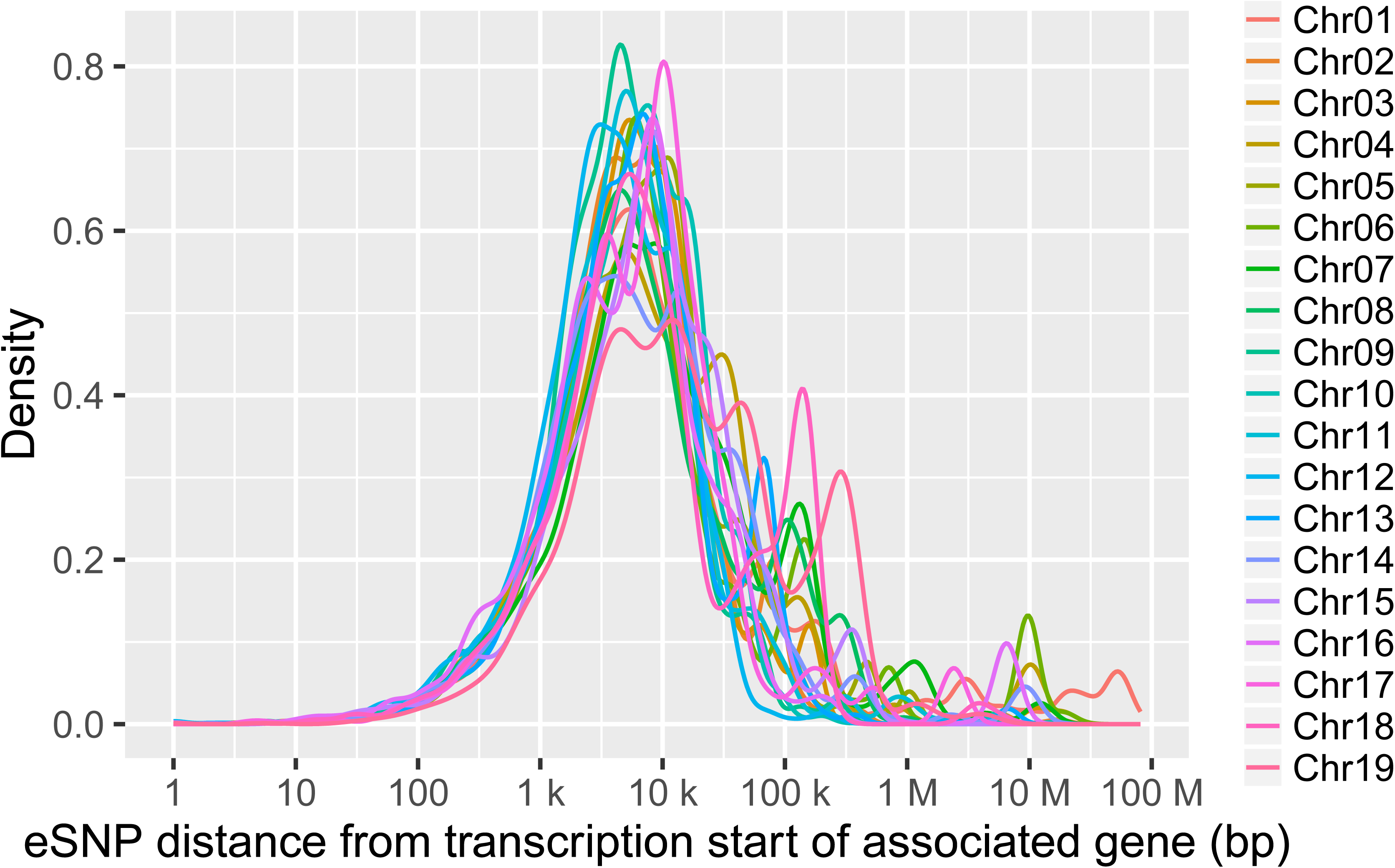
Physical distance between eSNPs and their associated genes for each of the 19 chromosomes.

**S5 Fig.**
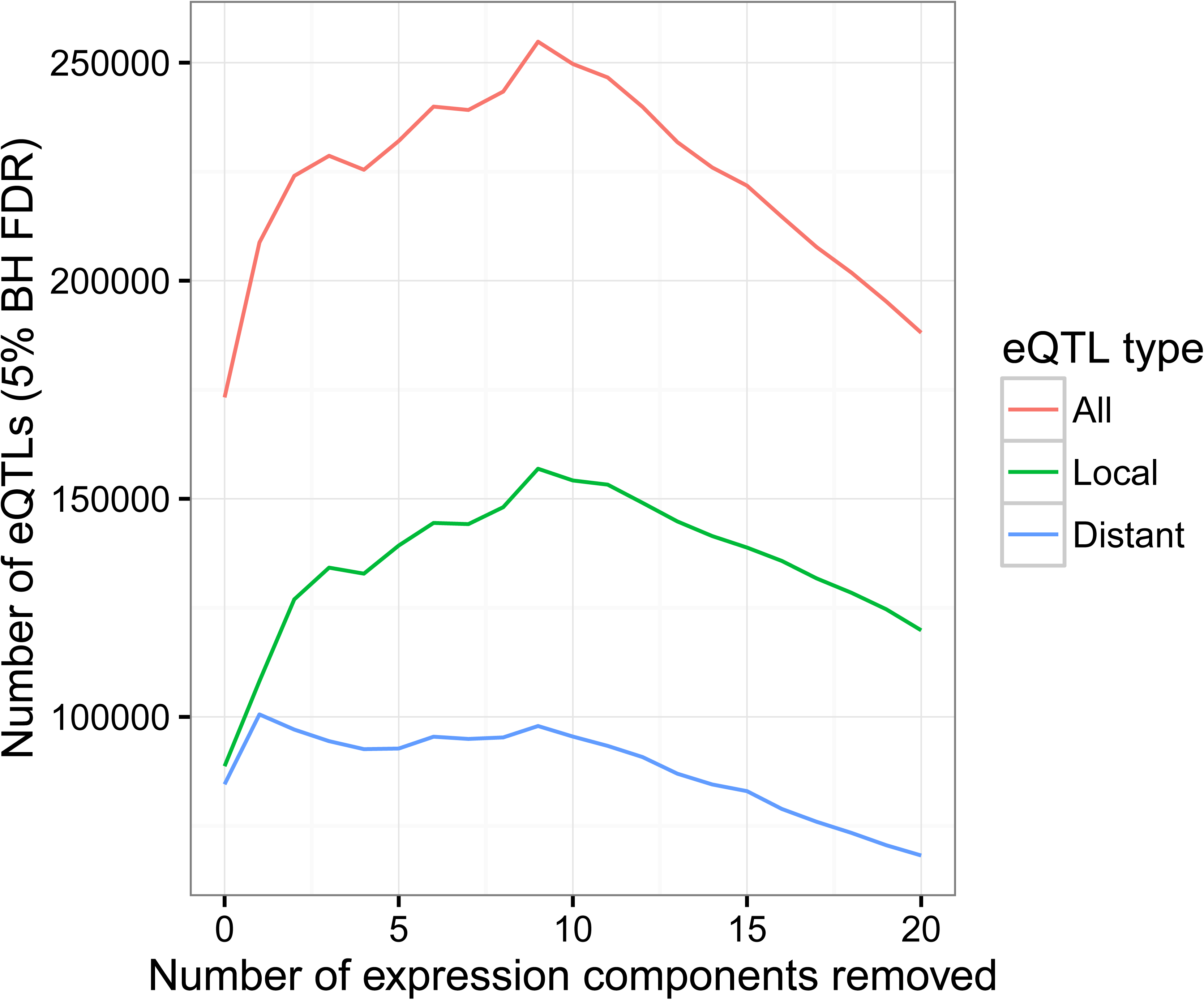
The number of eQTLs detected for different levels of hidden confounder removal. The y-axis shows the number of eQTLs detected at 5% FDR (prior to the empirical FDR calculation). The x-axis shows the number of principal components regressed out of the gene expression data. Distant eQTLs are defined as eQTLs where the gene and the SNP are located on different scaffolds.

**S6 Fig.**
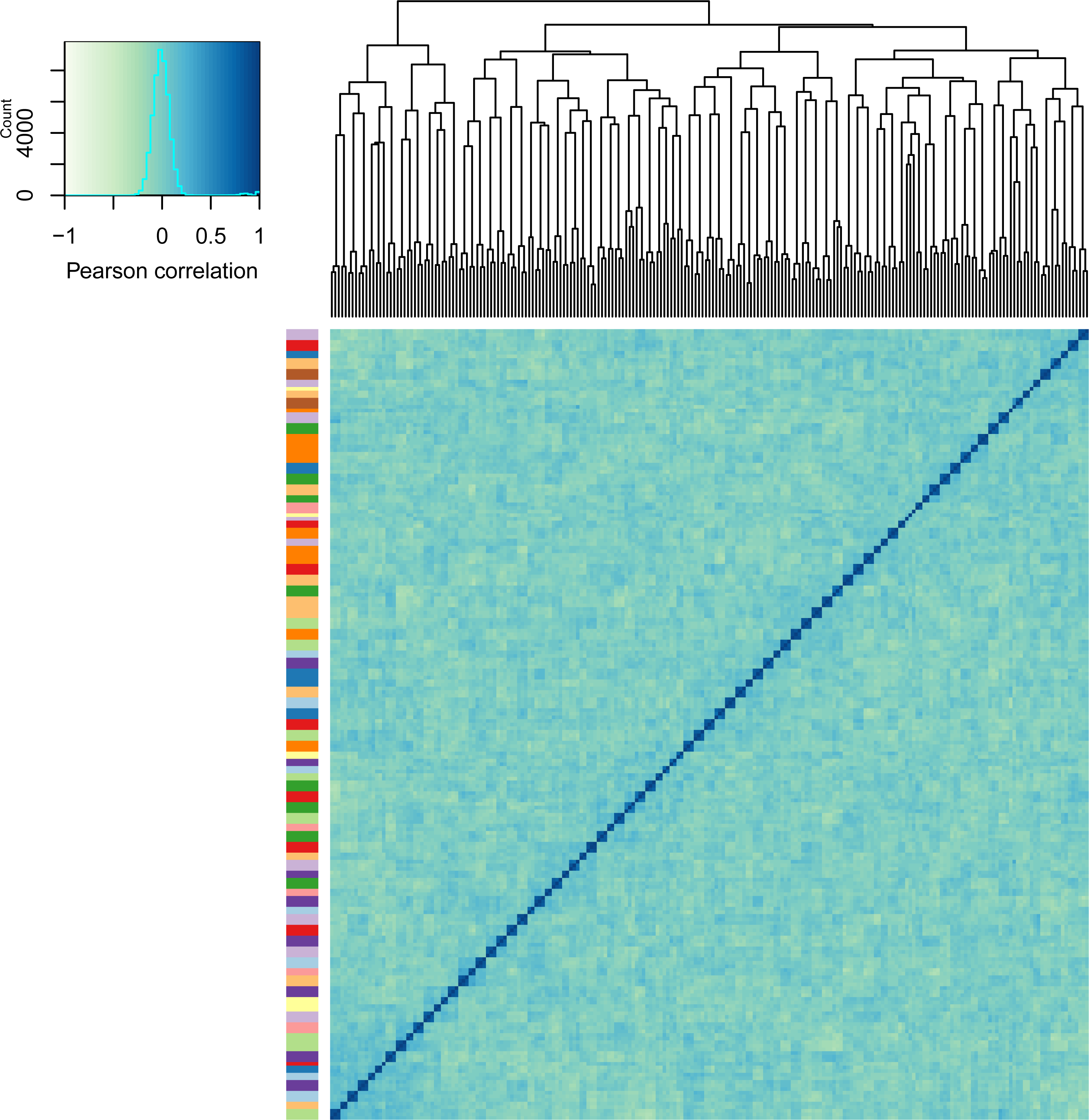
Heatmap representing the sample clustering based on gene expression values after hidden confounder removal. The 500 most variable genes in the original expression data were used for calculating the sample correlations (i.e. the same genes as in Fig 1D). The colour bar represents the population of the samples with the same colour scheme as in Fig 1D. The small clusters on the diagonal represent genotype replicates.

**S7 Fig.**
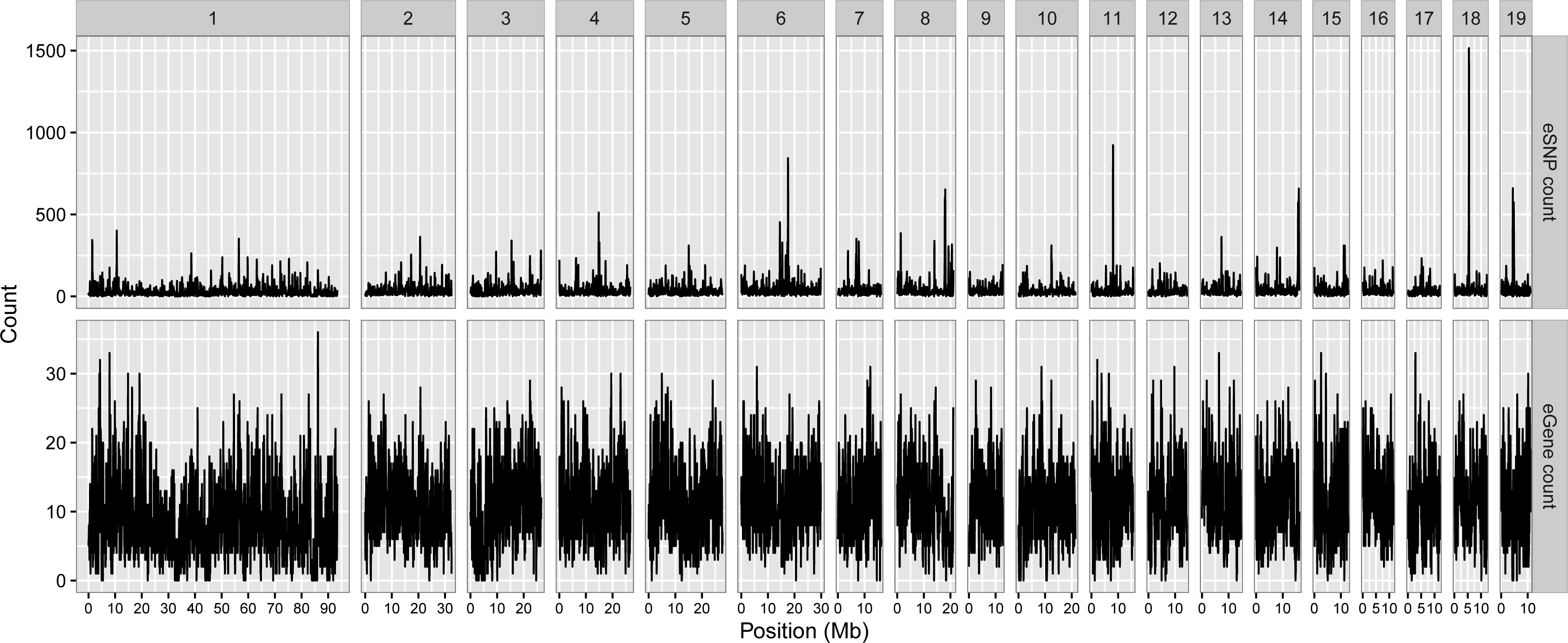
Sliding window approach to detect eQTL hotspots. The upper panel shows the number of eSNPs in genomic windows of 100 kb for each of the 19 chromosomes. The lower panel shows the number of unique genes that are associated to each genomic window (nominal eQTL p-value < 1×10^-6^).

**S8 Fig.**
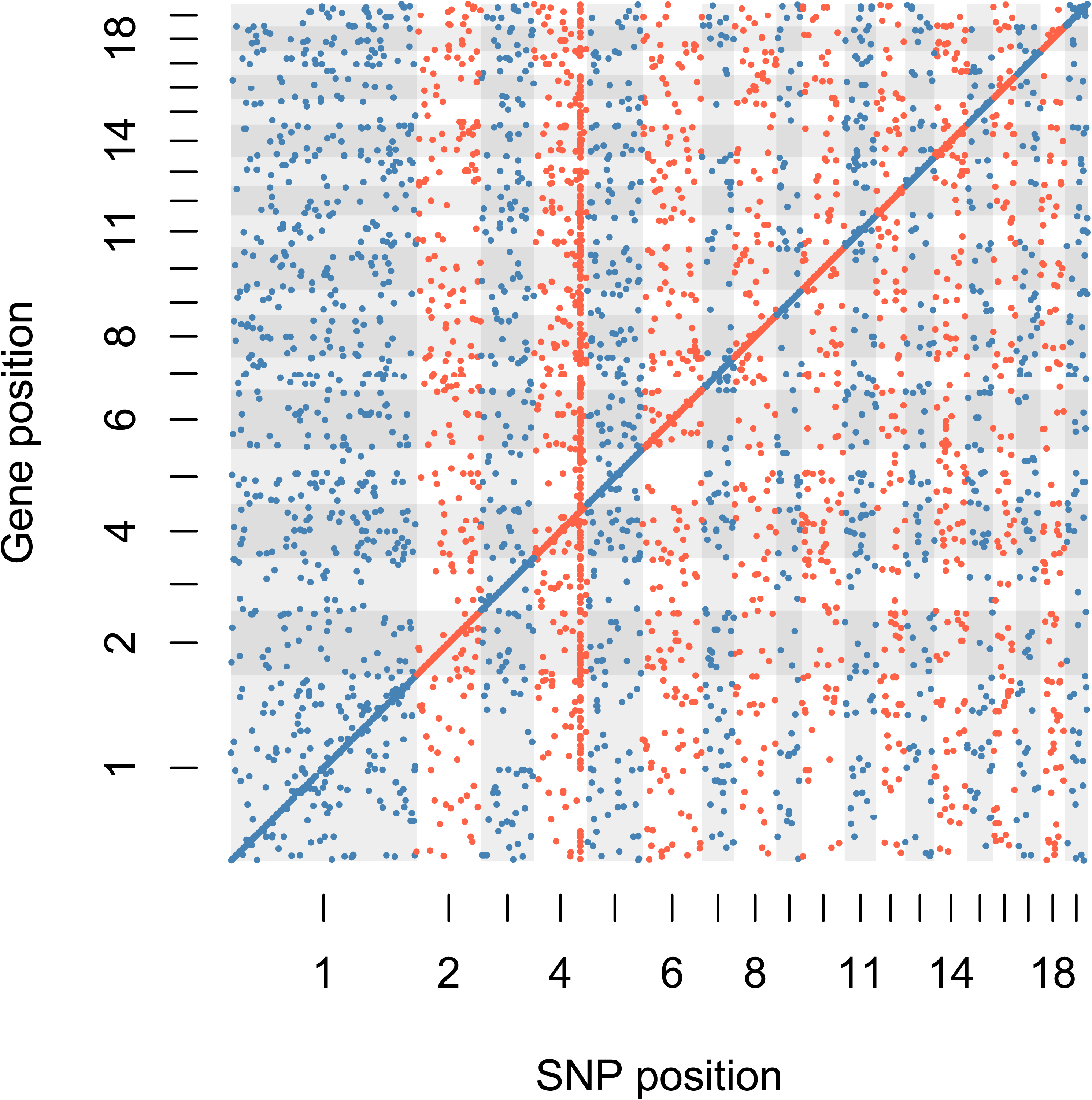
**eQTL scatter plot using gene expression data before hidden confounder removal.** A clear hotspot can be seen at the end of chromosome 4.

**S9 Fig.**
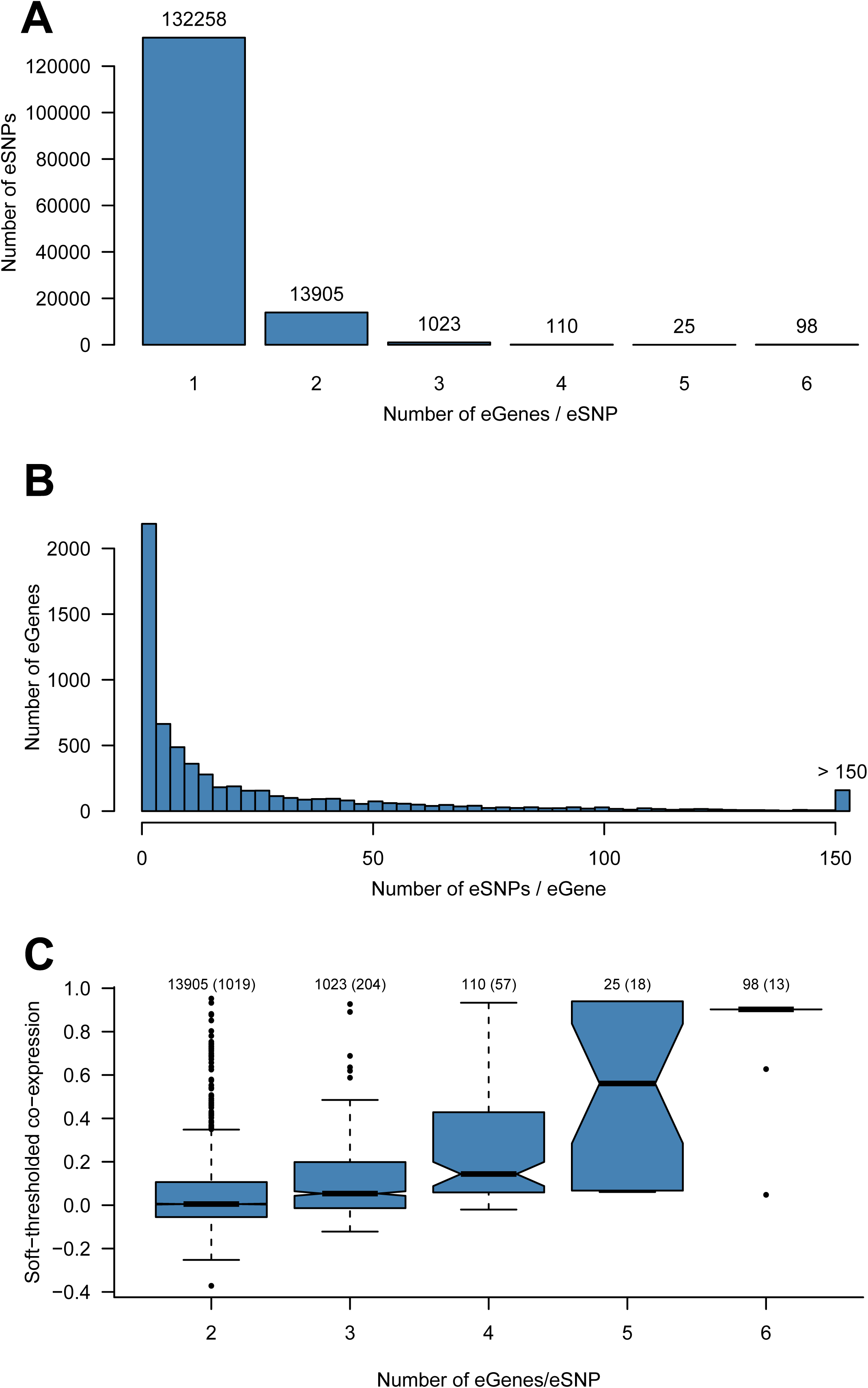
eSNPs per eGene and eGenes per eSNP. (A) Bar plot of the number of associated eGenes per eSNP. (B) Histogram of the number of associated eSNPs per eGene. (C) The mean co-expression for genes associated with the same eSNP. The numbers on the x-axis represents the number of eGenes that a single eSNP is associated to, and the numbers above the boxes shows the total number of eSNPs that are represented, and in parentheses are the number of unique genes that they are associated with.

**S10 Fig.**
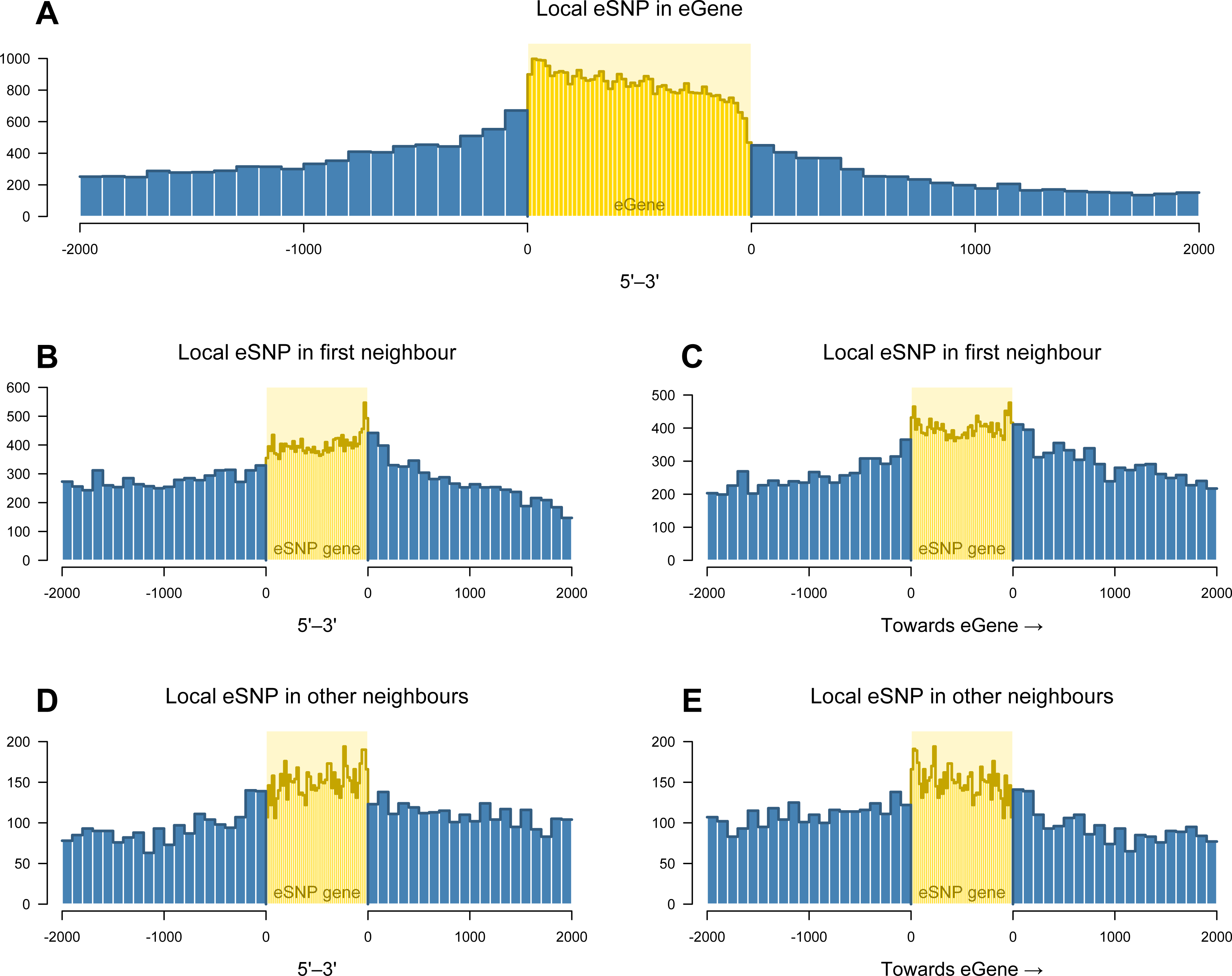
Local eSNP distribution. Local eSNPs were counted in cases where the eSNP was located in the gene it was associated with (A) as well as when it was located in a direct neighbour of the eGene (B, C) or in a gene further away from the eGene (D, E). For cases when the eSNP was located in neighbouring genes (B-E), this was plotted both in the direction of the gene (B, D) and relative to the eGene (C, E). Up and downstream regions were divided into 1 kbp bins while the intergenic region was normalised to 20 bins. In other words, up and downstream counts are not directly comparable to intragenic counts.

**S11 Fig.**
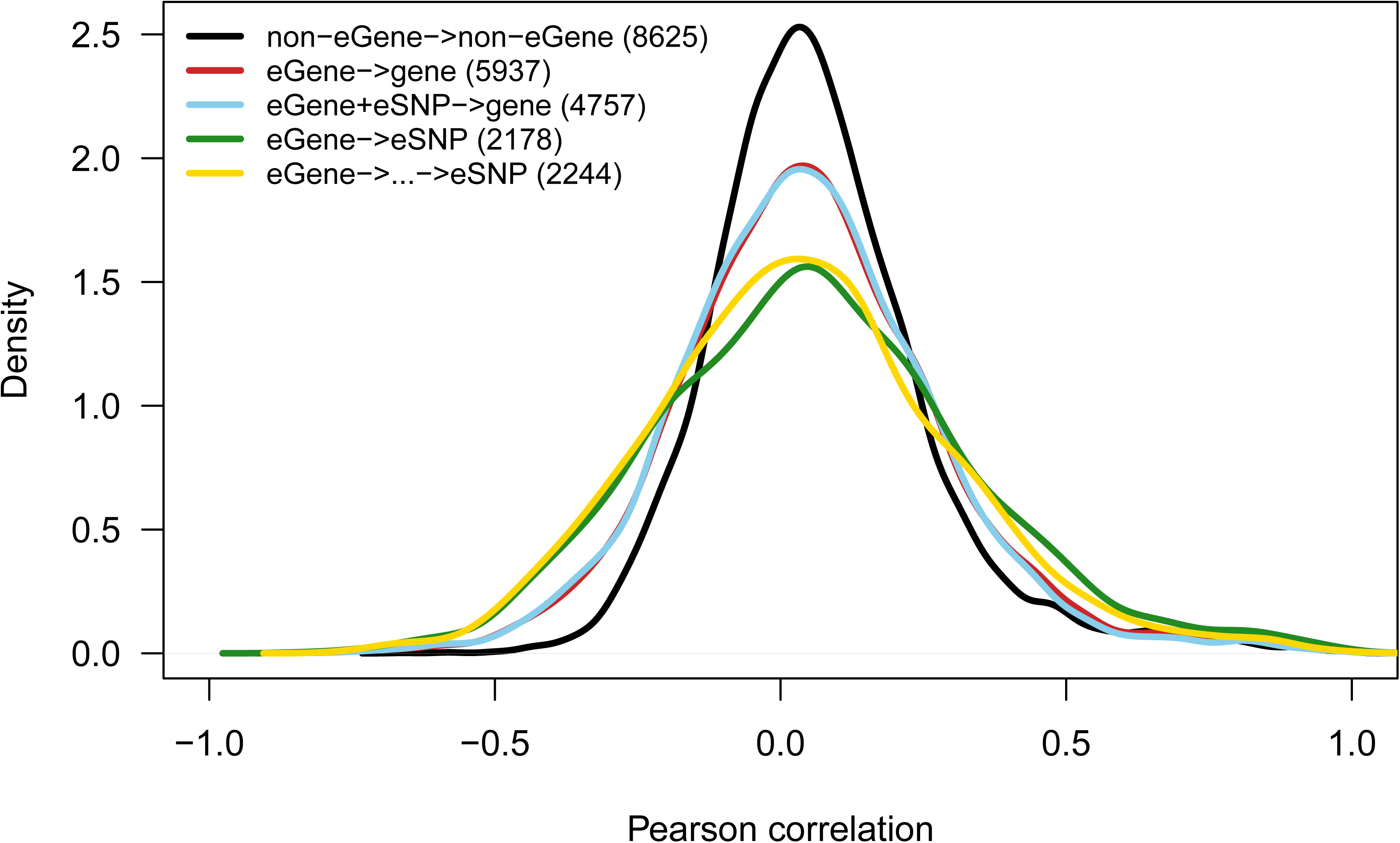
Gene expression correlation distributions for neighbouring genes. Neighbouring genes in the genome were divided into five different categories: non-eGene – non-eGene, eGene – gene, eGene with eSNP proximal to it - gene, eGene – first neighbour harbouring the eSNP, and eGene – neighbour further away harbouring the eSNP. The numbers in parenthesis in the legend represent the number of pairs in each category.

**S12 Fig.**
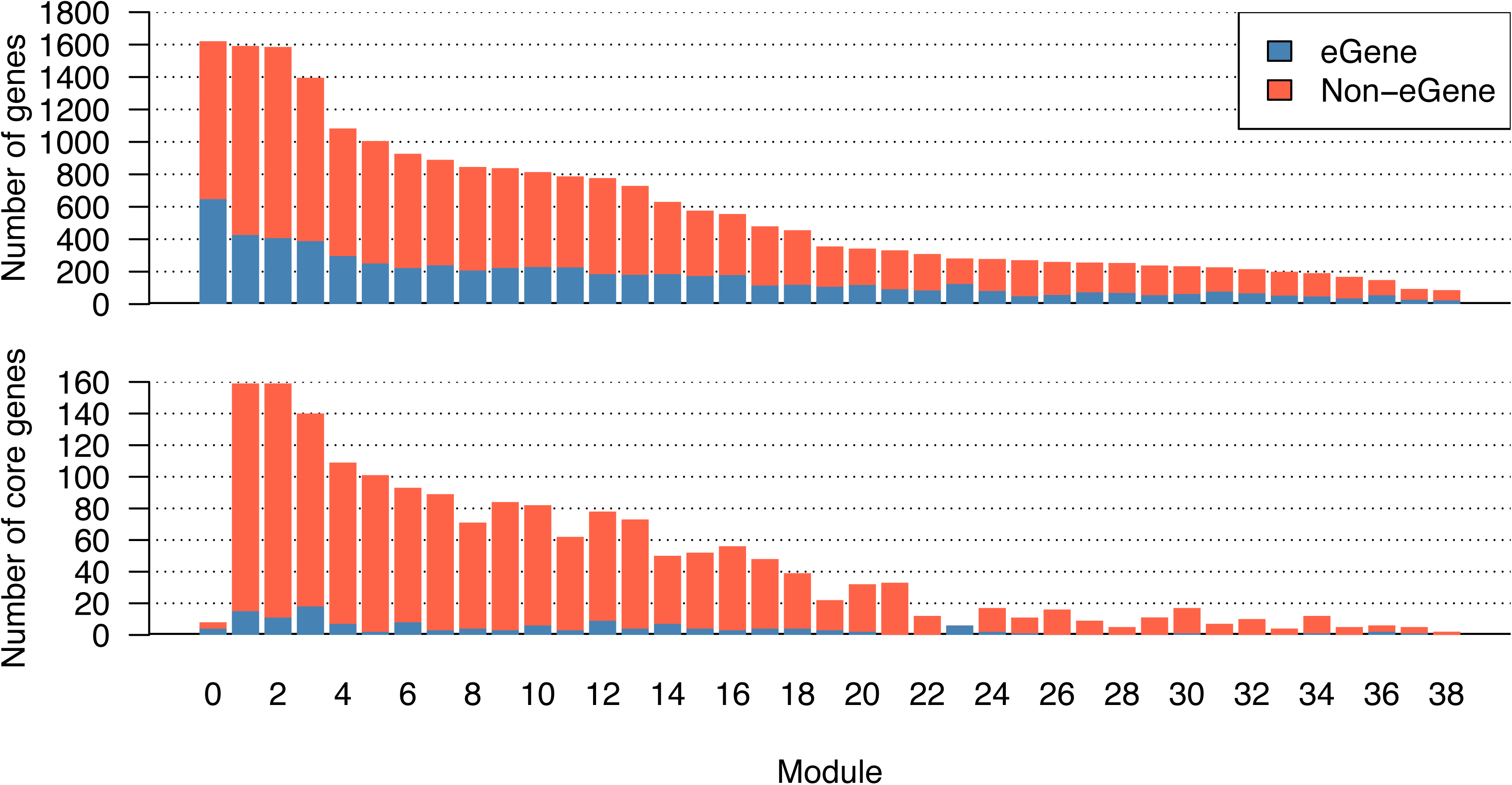
**The number of genes and the number of core genes in each network module.** Each bar is divided into eGene and non-eGene assignments. Module 0 contains genes not assigned to any module.

**S13 Fig.**
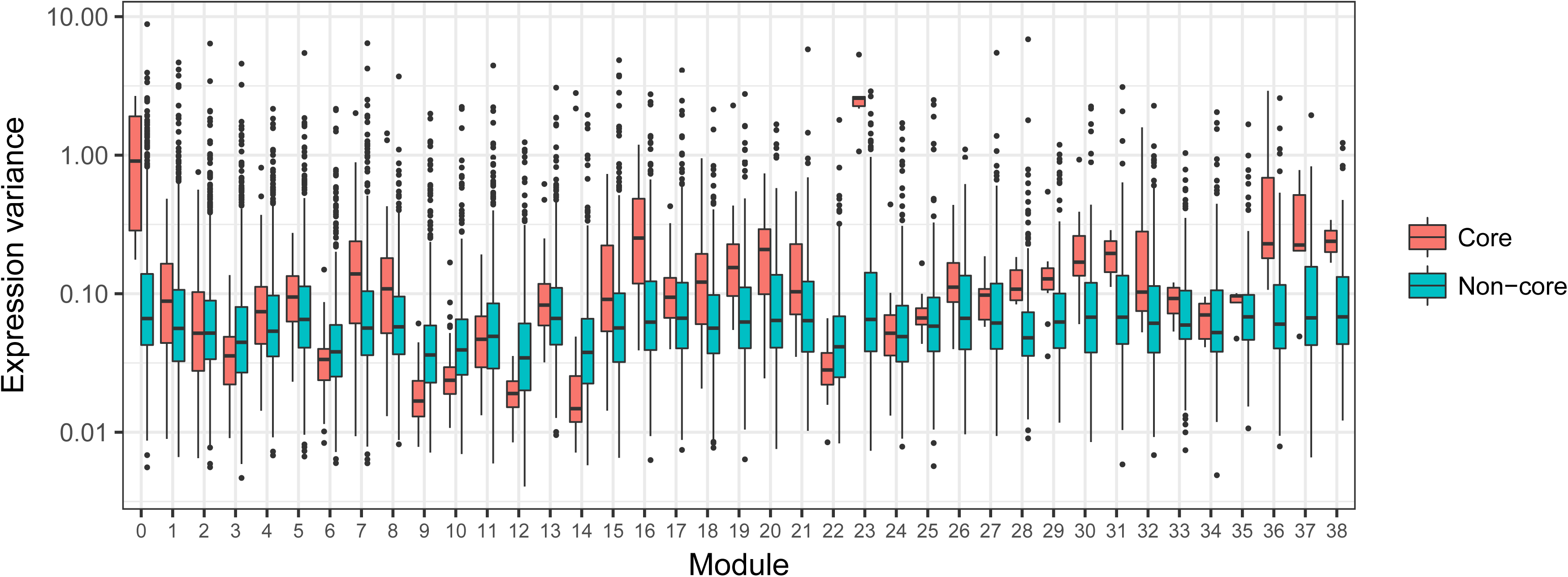
Gene expression variance in module cores and non-cores for each of the 38 modules.

**S14 Fig.**
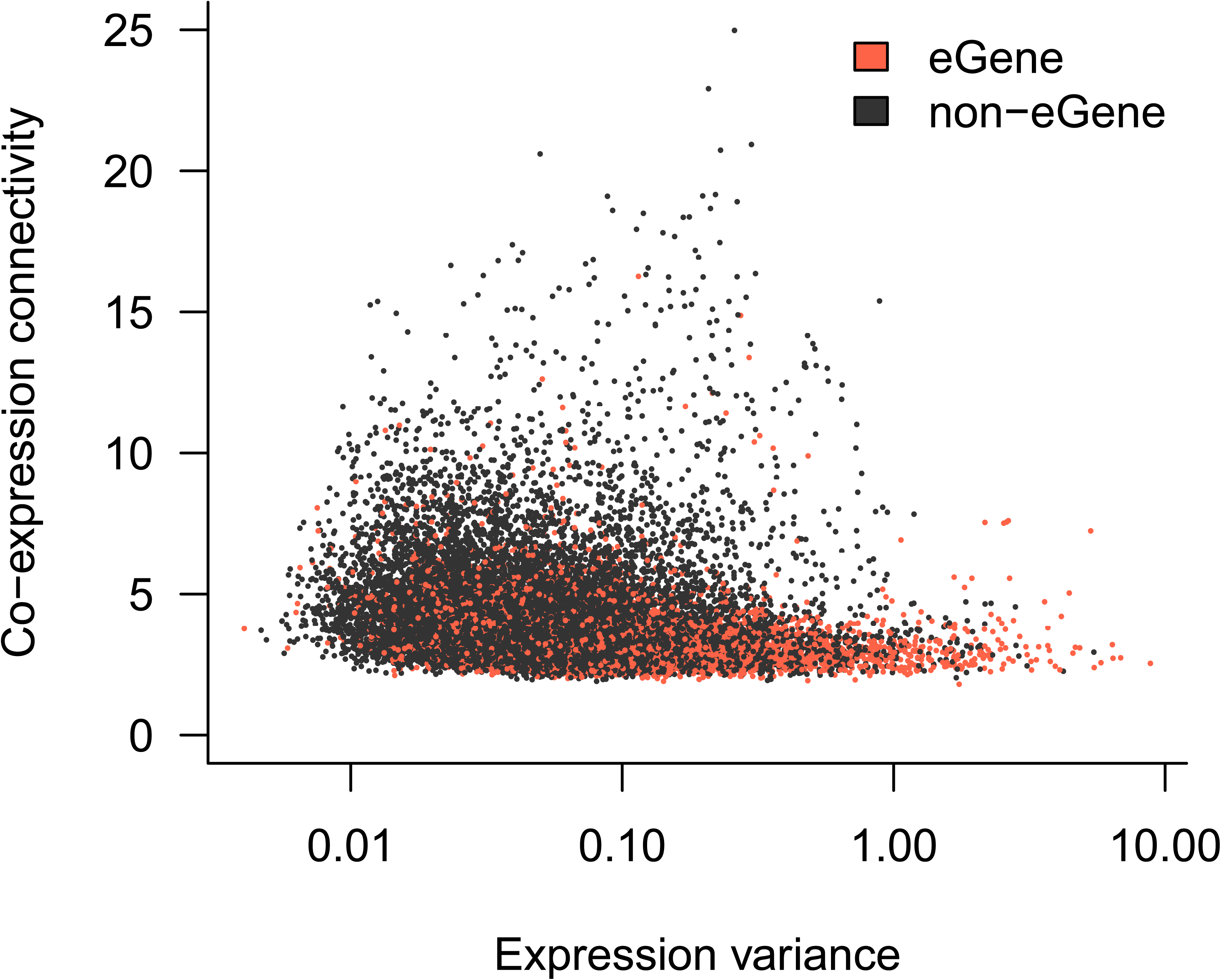
Gene expression variance plotted against co-expression network connectivity. eGenes are indicated by red points.

**S15 Fig.**
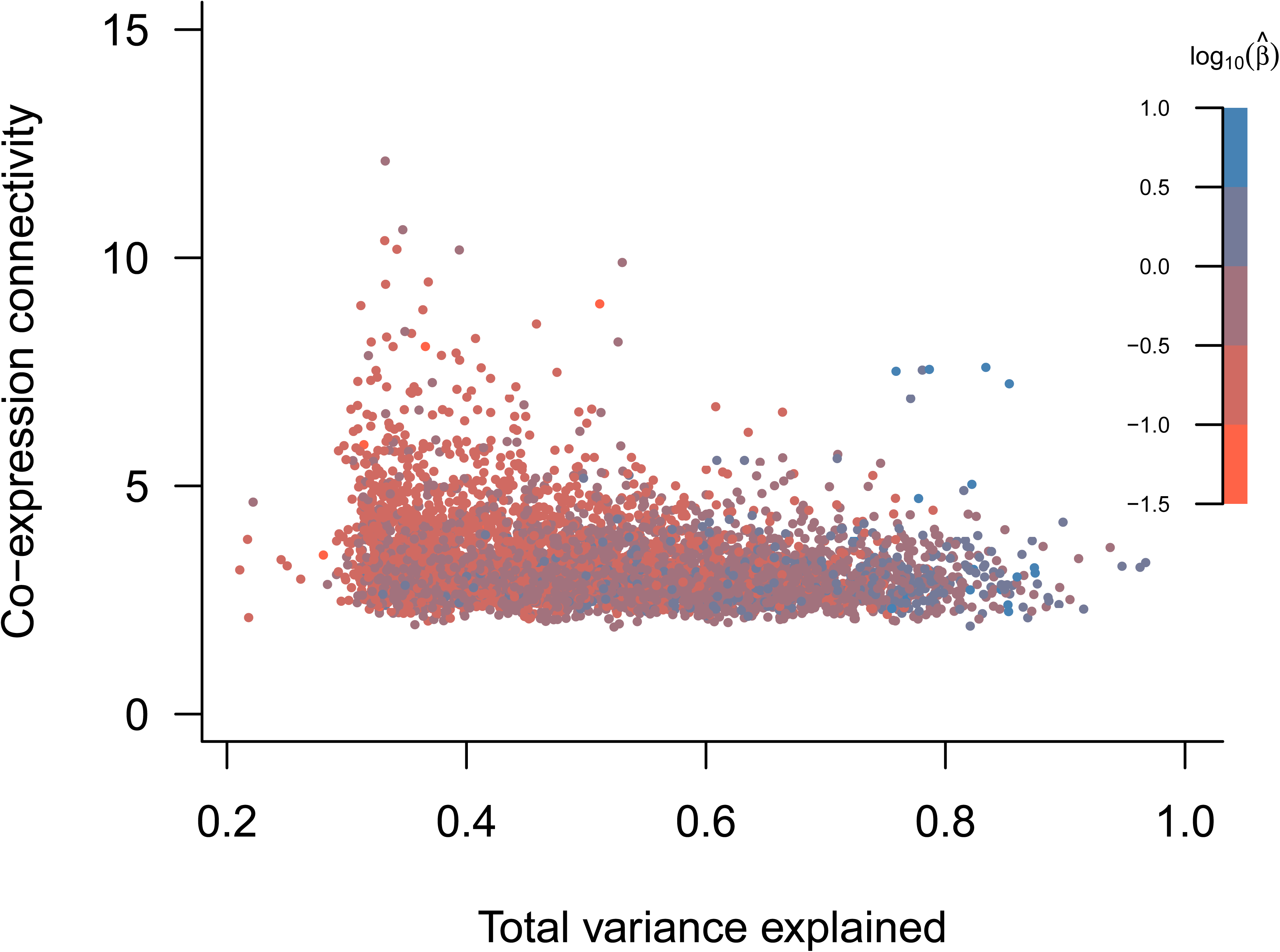
Relationship between total variance explained by eQTL and co-expression connectivity. The variance explained is the adjusted variance explained based on all eQTL associated with each gene.

**S16 Fig.**
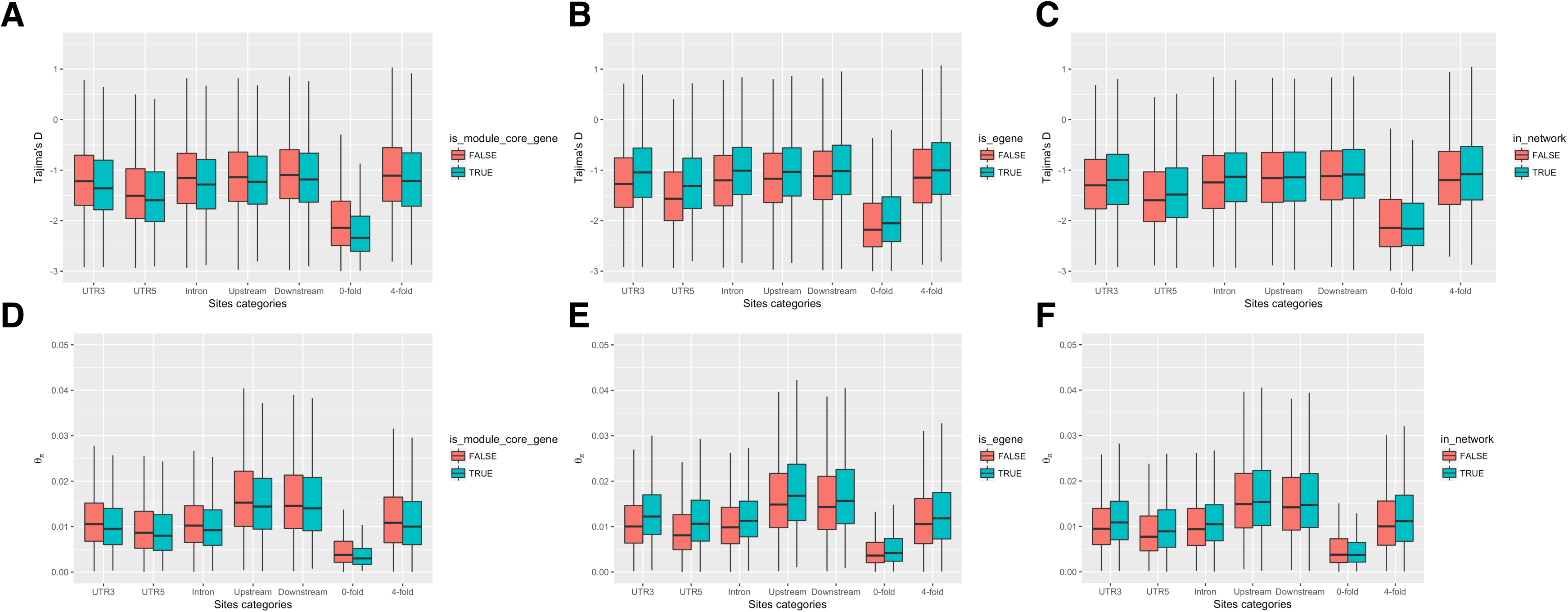
Comparisons of nucleotide diversity and Tajima’s D for different genomic features. Upstream and downstream are 1 kbp away from the gene start and end, respectively. 0-fold and 4- fold refers to 0-fold and 4-fold degenerate sites, respectively.

**S17 Fig.**
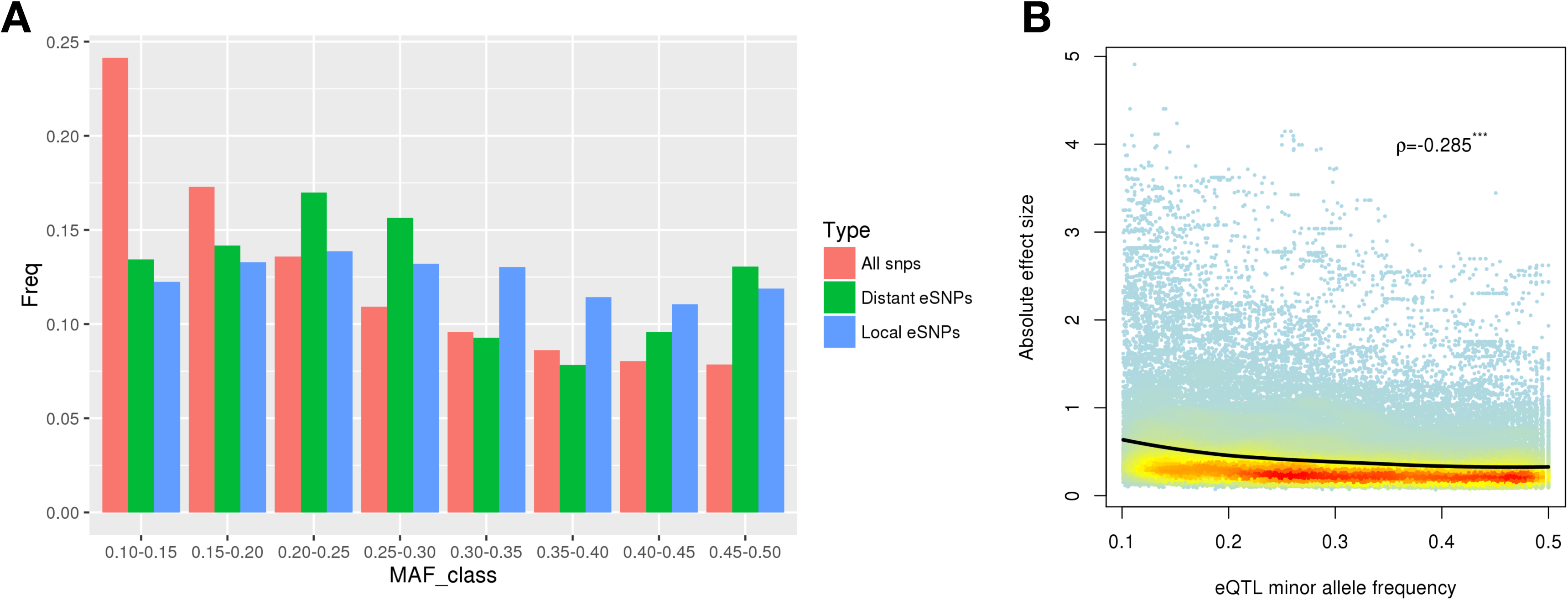
Minor allele frequency of eSNPs. (A) Comparison of minor allele frequency between all SNPs, distant eSNPs and local eSNPs. (B) The relationship between minor allele frequency and effect size (absolute value of beta) of eQTLs.

**S18 Fig.**
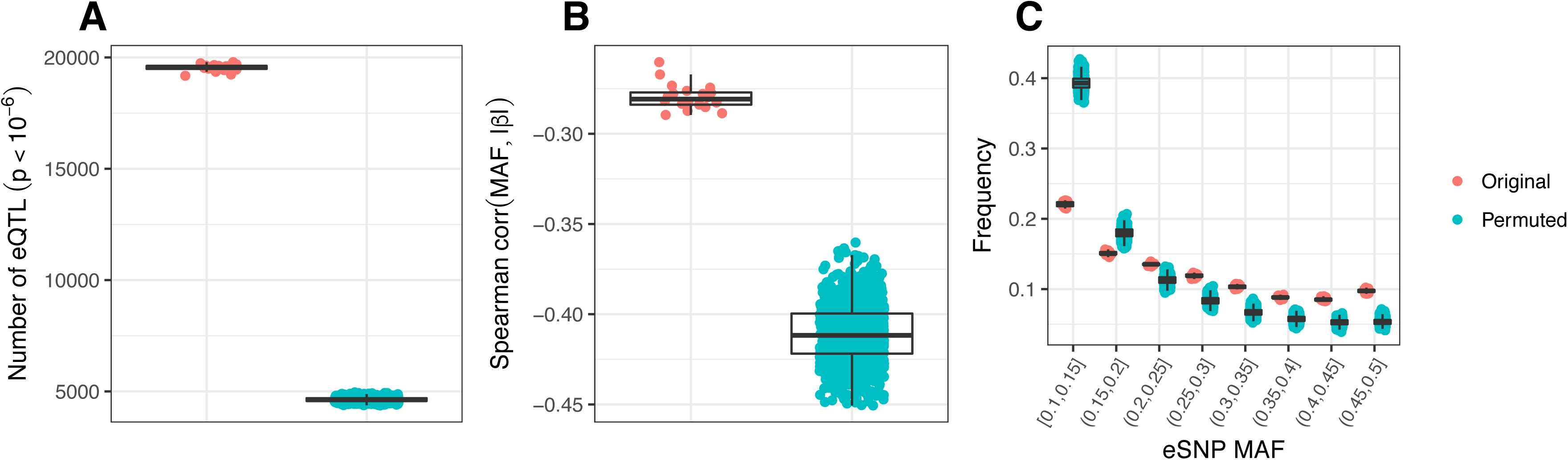
Relationship between MAF and effect size in permuted eQTLs. eQTL permutations were done by selecting 20 random subsets of 150,000 SNPs each. Each of these sets were associated with gene expression levels, and in addition, 50 permutations were performed for each set by shuffling the genotype labels of the SNP data, resulting in a total of 1,000 permuted eQTL sets. (A) The number of SNPs for the nominal p-value threshold 10^Λ^-6 (the same initial threshold that was used for the original associations). (B) The Spearman correlation between minor allele frequency and absolute effect size for the 20 sets and their corresponding permutations. (C) Allele frequency spectra for the 20 subsets and their permutations.

**S19 Fig.**
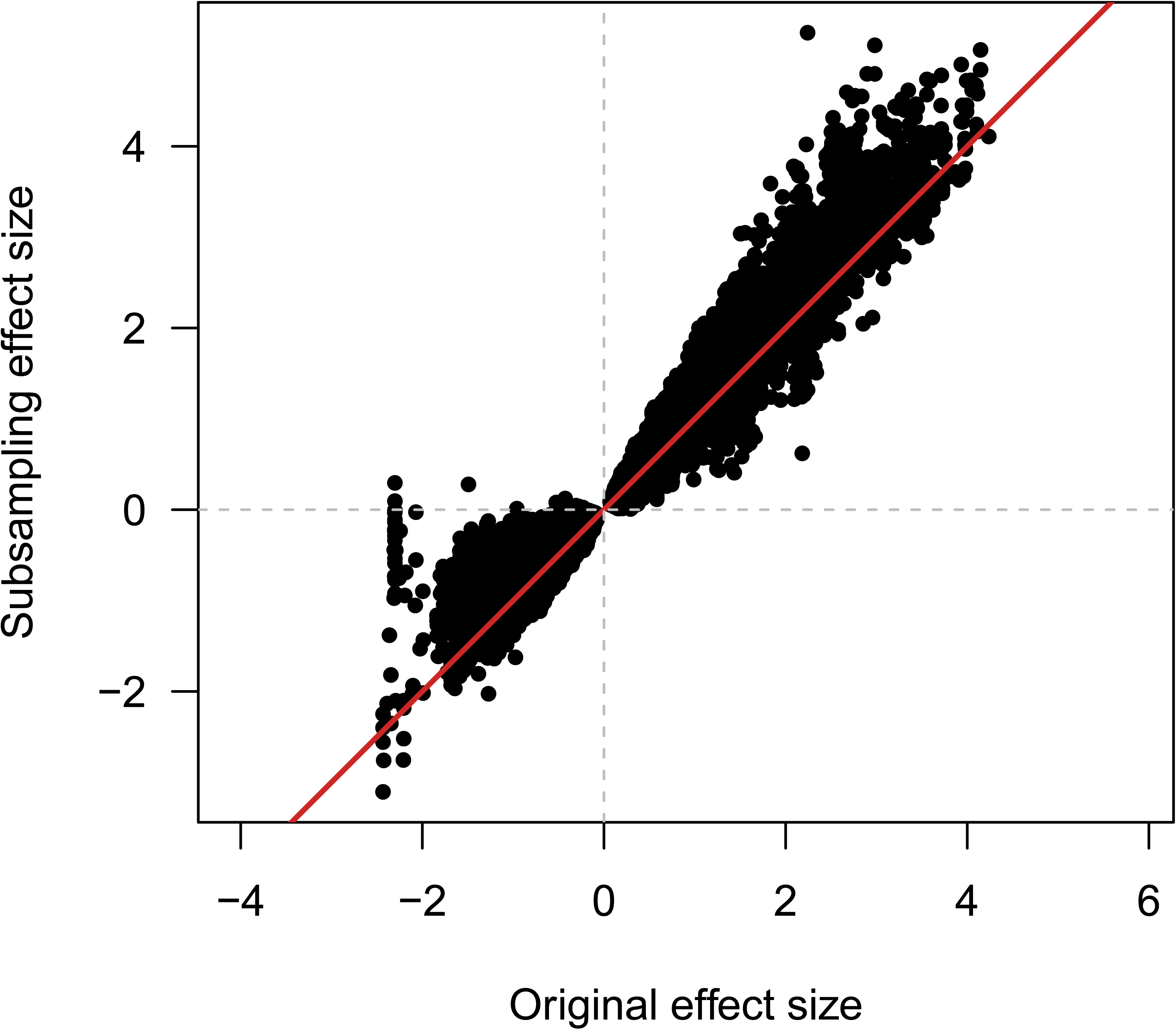
Comparing the effect sizes between the original and the subsampled data. The Pearson correlation is 0.98 (df = 157,020).

**S20 Fig.**
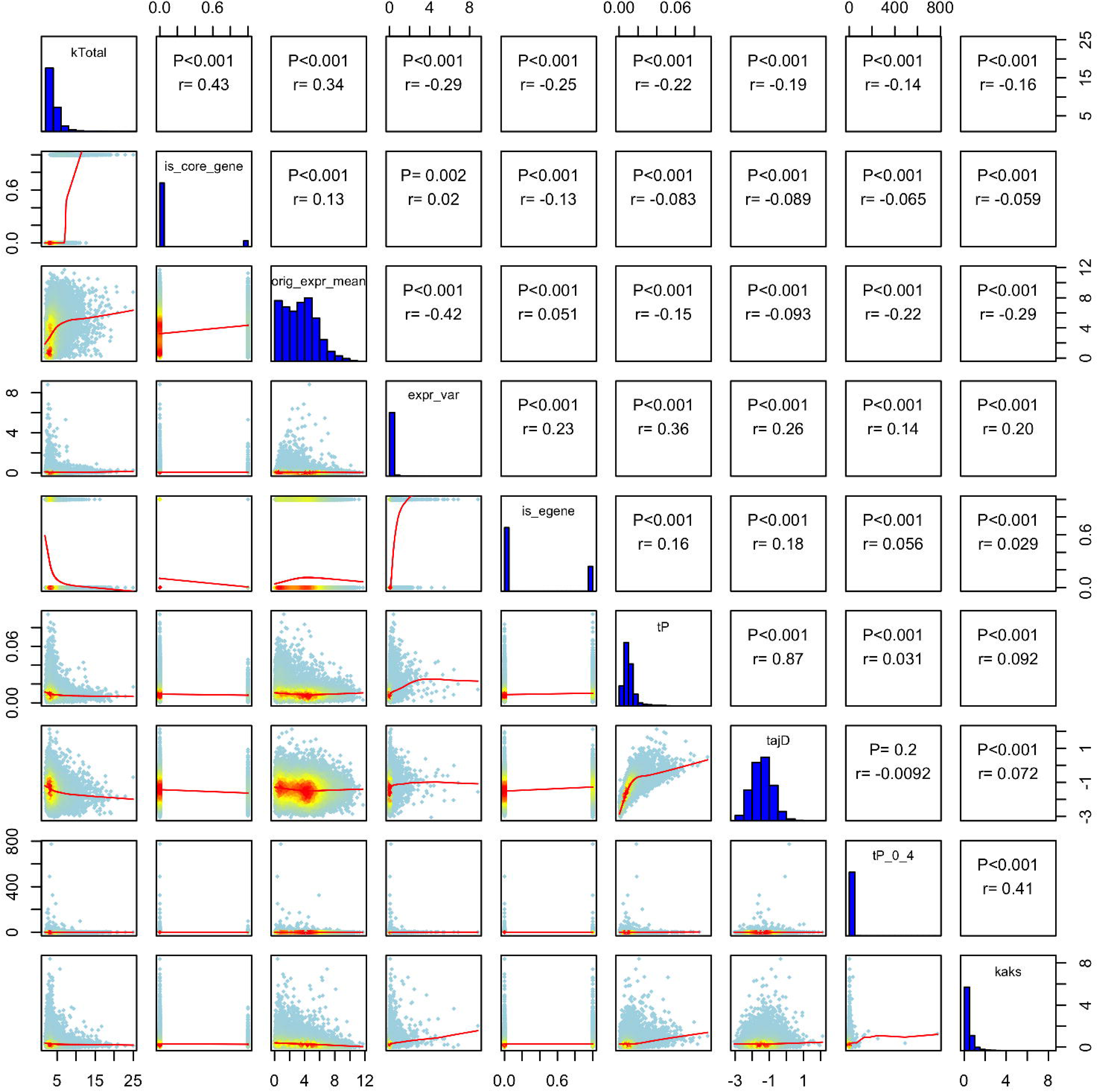
Association between metrics of gene expression and sequence evolution. Scatter plots (lower off-diagnal, the red-to-yellow-to-blue gradient indicates decreased density of observed events at a give location in the graph) and correlations with probability values (upper off-diagonal, spearman’s rank correlations) for measures of gene expression and sequence evolution: kTotal, the total (global) connectivity in the network; is_core_gene, gene is part of module core or not; orig_expr_mean, mean expression before hidden confounder removal; orig_expr_var, expression variance before hidden confounder removal; is_egene, gene with eQTLs or not; tP, pairwise nucleotide diversity; tajD, Tajima’s D; tP_0_4, the ratio of pairwise nucleotide diversity at zero-fold non-synonymous and four-fold synonymous sites; kaks, dN/dS.

**S21 Fig.**
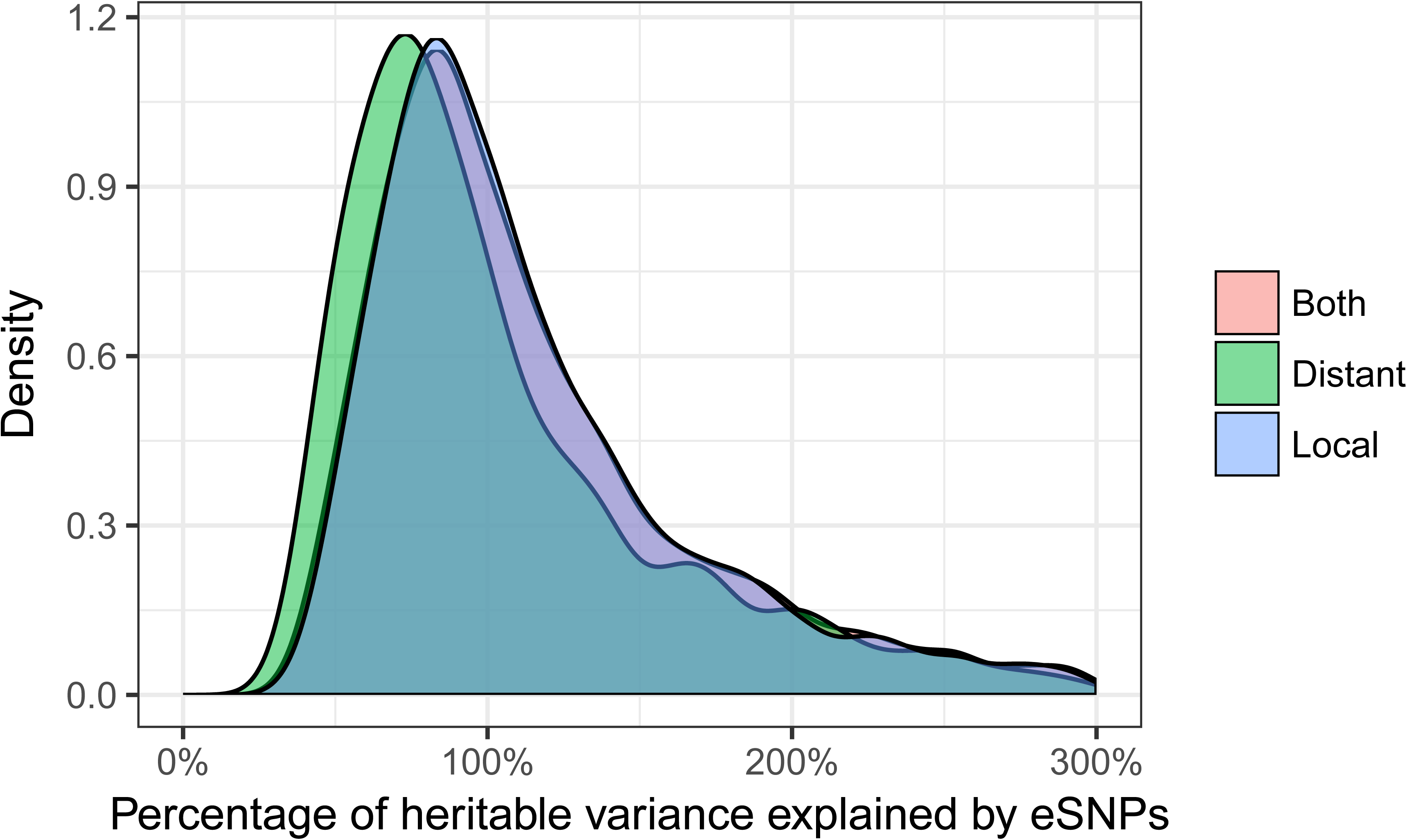
Variance explained by eSNPs and heritability. This figure assumes that eSNPs only account for heritable variation, explaining why some of the eSNPs explain more than 100% of the heritability. The x-axis has been truncated at 300%.

**S22 Fig.**
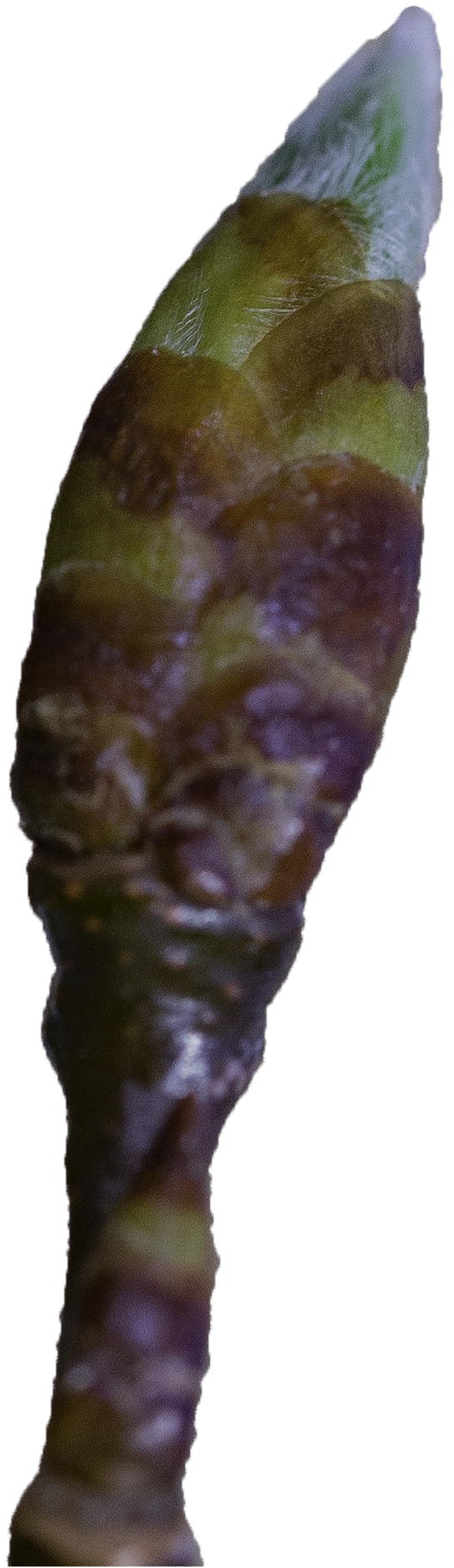
Representative photo of the sampled bud flush stage.

**S23 Fig.**
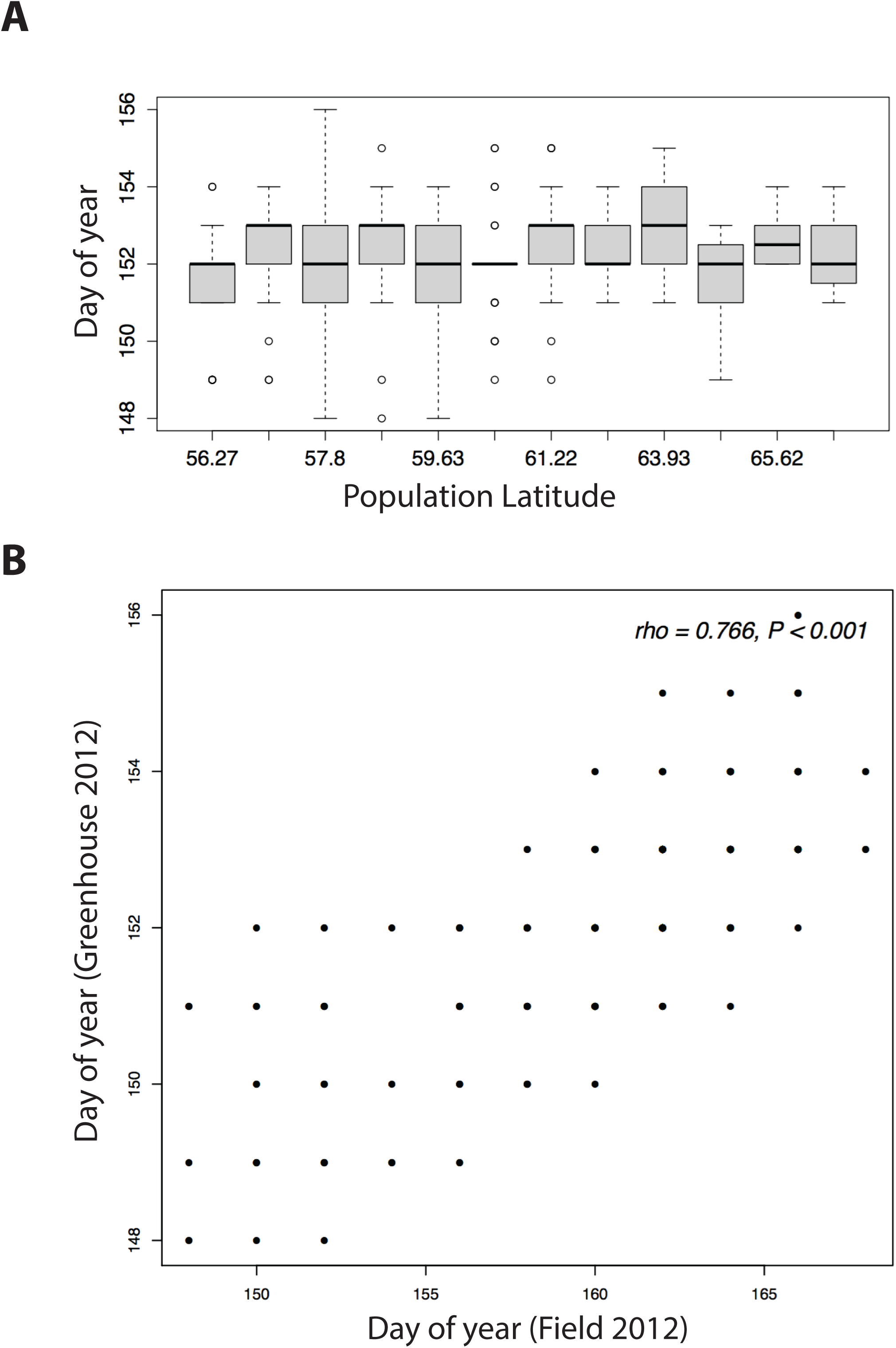
Sampling time and day of bud flush. (A) Box plot distributions of the Julian day of sampling for the SwAsp sub-populations. (B) The relationship between Julian day of bud flush for the greenhouse sampled buds and Julian day of bud flush in the field for the same year (2012).

**S24 Fig.**
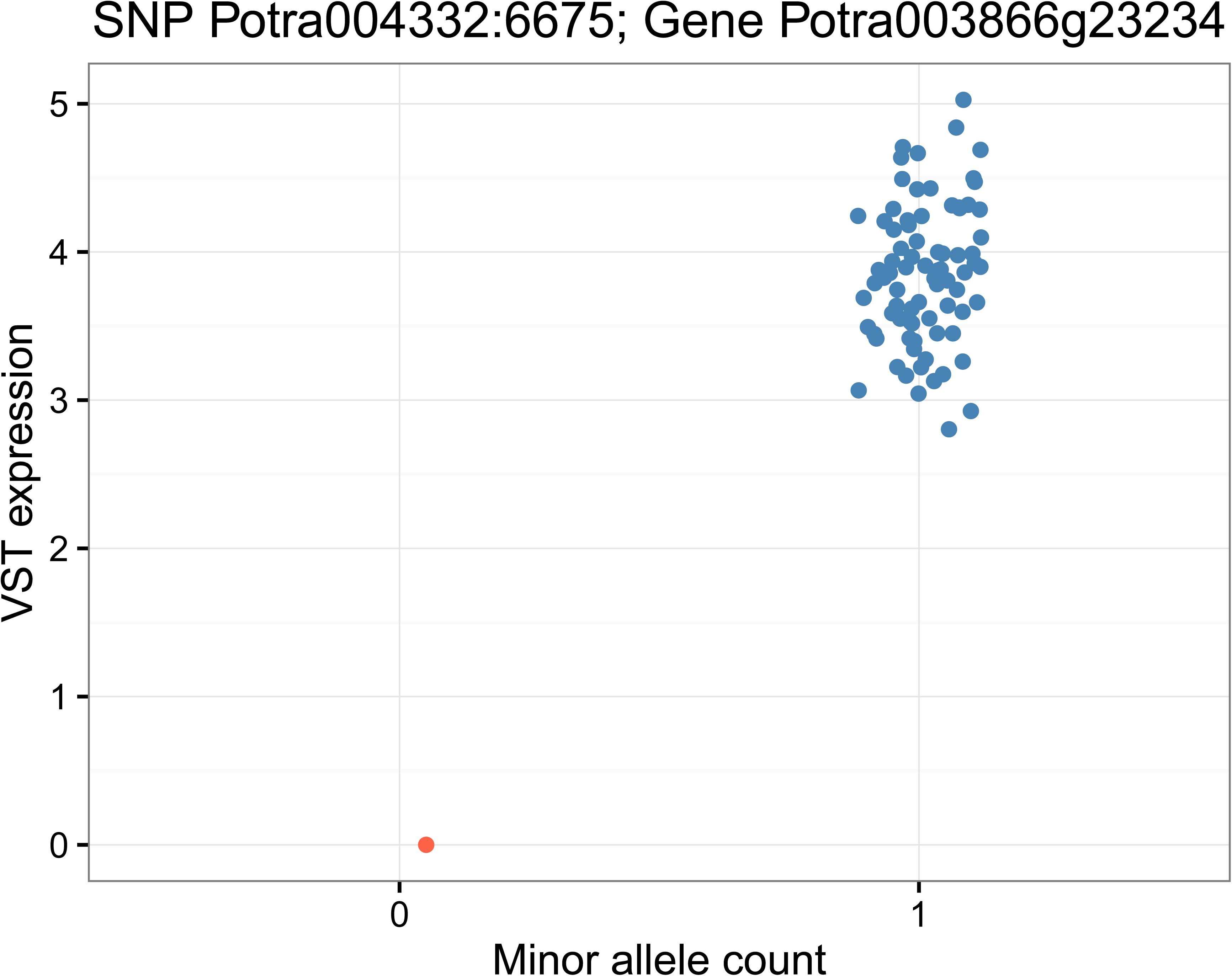
Motivating example for major genotype filtering. In this case the SNP is heterozygous in all samples but one, resulting in a minor allele frequency close to 0.5, but a major genotype frequency close to one. The resulting association turns out very significant, but is only supported by a single sample.

**S25 Fig.**
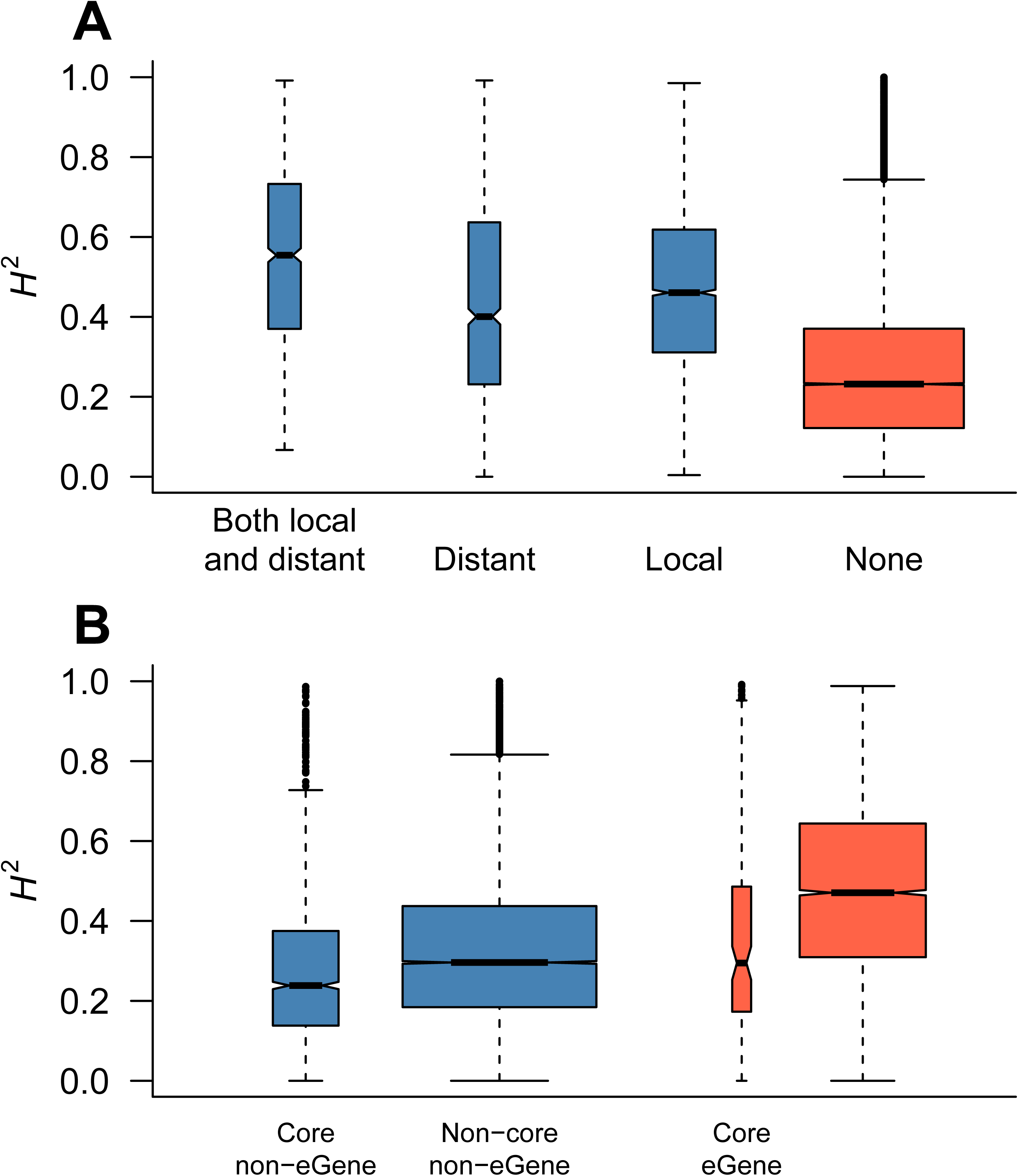
**Heritability distributions for four non-overlapping sets of genes:** genes with both local and distant eQTLs, genes with only distant eQTLs, genes with only local eQTLs, and genes with no significant eQTLs. The box widths are proportional to the number of genes in each set.

## References

1. Ingvarsson PK, Hvidsten TR, Street NR. Towards integration of population and comparative genomics in forest trees. New Phytol. 2016; doi:10.1111/nph.14153

2. Sandberg R, Yasuda R, Pankratz DG, Carter TA, Del Rio JA, Wodicka L, et al. Regional and strain-specific gene expression mapping in the adult mouse brain. Proc Natl Acad Sci U S A. 2000;97: 11038–43. doi:10.1073/pnas.97.20.11038

3. Primig M, Williams RM, Winzeler EA, Tevzadze GG, Conway AR, Hwang SY, et al. The core meiotic transcriptome in budding yeasts. Nat Genet. 2000;26: 415–423. doi:10.1038/82539

4. Jin W, Riley RM, Wolfinger RD, White KP, Passador-Gurgel G, Gibson G. The contributions of sex, genotype and age to transcriptional variance in Drosophila melanogaster. Nat Genet. 2001;29: 389–95. doi:10.1038/ng766

5. Oleksiak MF, Churchill GA, Crawford DL. Variation in gene expression within and among natural populations. Nat Genet. 2002;32: 261–266. doi:10.1038/ng983

6. Schadt EE, Monks SA, Drake TA, Lusis AJ, Che N, Colinayo V, et al. Genetics of gene expression surveyed in maize, mouse and man. Nature. 2003;422: 297–302. doi:10.1038/nature01434

7. Kirst M, Myburg AA, De León JPG, Kirst ME, Scott J, Sederoff R. Coordinated genetic regulation of growth and lignin revealed by quantitative trait locus analysis of cDNA microarray data in an interspecific backcross of eucalyptus. Plant Physiol. 2004;135: 2368–78. doi:10.1104/pp.103.037960

8. Morley M, Molony CM, Weber TM, Devlin JL, Ewens KG, Spielman RS, et al. Genetic analysis of genome-wide variation in human gene expression. Nature. 2004;430: 743–747. doi:10.1038/nature02797

9. Cheung VG, Spielman RS, Ewens KG, Weber TM, Morley M, Burdick JT. Mapping determinants of human gene expression by regional and genome-wide association. Nature. 2005;437: 1365–9. doi:10.1038/nature04244

10. Hubner N, Wallace CA, Zimdahl H, Petretto E, Schulz H, Maciver F, et al. Integrated transcriptional profiling and linkage analysis for identification of genes underlying disease. Nat Genet. 2005;37: 243–253. doi:10.1038/ng1522

11. Stranger BE, Forrest MS, Clark AG, Minichiello MJ, Deutsch S, Lyle R, et al. Genome-wide associations of gene expression variation in humans. PLoS Genet. 2005;1: 0695–0704. doi:10.1371/journal.pgen.0010078

12. DeCook R, Lall S, Nettleton D, Howell SH. Genetic Regulation of Gene Expression During Shoot Development in Arabidopsis. Genetics. 2006;172: 1155–1164. Available: http://www.genetics.org/cgi/content/abstract/172/2/1155

13. Dixon AL, Liang L, Moffatt MF, Chen W, Heath S, Wong KCC, et al. A genome-wide association study of global gene expression. Nat Genet. 2007;39: 1202–1207. doi:10.1038/ng2109

14. Gilad Y, Rifkin SA, Pritchard JK. Revealing the architecture of gene regulation: the promise of eQTL studies. Trends Genet. 2008;24: 408–415. doi:10.1016/j.tig.2008.06.001

15. Kim J, Gibson G. Insights from GWAS into the quantitative genetics of transcription in humans. Genet Res (Camb). 2010;92: 361–9. doi:10.1017/S001667231000056X

16. Powell JE, Henders AK, McRae AF, Kim J, Hemani G, Martin NG, et al. Congruence of Additive and Non-Additive Effects on Gene Expression Estimated from Pedigree and SNP Data. PLoS Genet. 2013;9. doi:10.1371/journal.pgen.1003502

17. Wang RL, Stec A, Hey J, Lukens L, Doebley J. The limits of selection during maize domestication. Nature. 1999;398: 236–239. doi:10.1038/18435

18. Carroll SB. Endless forms: the evolution of gene regulation and morphological diversity. Cell. 2000;101: 577–580. doi:10.1016/S0092-8674(00)80868-5

19. Brem RB, Yvert G, Clinton R, Kruglyak L. Genetic dissection of transcriptional regulation in budding yeast. Science (80- ). 2002;296: 752–5. doi:10.1126/science.1069516

20. Ayroles JF, Carbone MA, Stone E a, Jordan KW, Lyman RF, Magwire MM, et al. Systems genetics of complex traits in Drosophila melanogaster. Nat Genet. 2009;41: 299–307. doi:10.1038/ng.332

21. Mackay TFC, Stone EA, Ayroles JF. The genetics of quantitative traits: challenges and prospects. Nat Rev Genet. 2009;10: 565–77. doi:10.1038/nrg2612

22. Liao B-Y, Weng M-P, Zhang J. Contrasting genetic paths to morphological and physiological evolution. Proc Natl Acad Sci U S A. 2010;107: 7353–8. doi:10.1073/pnas.0910339107

23. Hines HM, Papa R, Ruiz M, Papanicolaou A, Wang C, Nijhout HF, et al. Transcriptome analysis reveals novel patterning and pigmentation genes underlying Heliconius butterfly wing pattern variation. BMC Genomics. 2012;13: 288. doi:10.1186/1471-2164-13-288

24. Richards CL, Rosas U, Banta J, Bhambhra N, Purugganan MD. Genome-Wide Patterns of Arabidopsis Gene Expression in Nature. PLoS Genet. 2012;8: e1002662. doi:10.1371/journal.pgen.1002662

25. Jansen RC, Nap JP. Genetical genomics: the added value from segregation. Trends Genet. 2001;17: 388–91. doi:10.1016/S0168-9525(01)02310-1

26. Doerge RW. Mapping and analysis of quantitative trait loci in experimental populations. Nat Rev Genet. 2002;3: 43–52. doi:10.1038/nrg703

27. Josephs EB, Lee YW, Stinchcombe JR, Wright SI. Association mapping reveals the role of purifying selection in the maintenance of genomic variation in gene expression. Proc Natl Acad Sci U S A. 2015;112: 15390–5. doi:10.1073/pnas.1503027112

28. Lappalainen T. Functional genomics bridges the gap between quantitative genetics and molecular biology. Genome Res. 2015;25: 1427–1431. doi:10.1101/gr.190983.115.

29. Flint J, and Mackay TFC. Genetic architecture of quantitative traits in flies, mice and humans. Genome Res. 2009;19: 723–733. doi:10.1101/gr.086660.108

30. Hindorff L a, Sethupathy P, Junkins H a, Ramos EM, Mehta JP, Collins FS, et al. Potential etiologic and functional implications of genome-wide association loci for human diseases and traits. Proc Natl Acad Sci U S A. 2009;106: 9362–7. doi:10.1073/pnas.0903103106

31. Ku CS, Loy EY, Pawitan Y, Chia KS. The pursuit of genome-wide association studies: where are we now? J Hum Genet. 2010;55: 195–206. doi:10.1038/jhg.2010.19

32. Wray GA. The evolutionary significance of cis-regulatory mutations. Nat Rev Genet. 2007;8: 206–16. doi:10.1038/nrg2063

33. Kliebenstein DJ, West MAL, van Leeuwen H, Kim K, Doerge RW, Michelmore RW, et al. Genomic survey of gene expression diversity in Arabidopsis thaliana. Genetics. 2006;172: 1179–89. doi:10.1534/genetics.105.049353

34. Keurentjes JJB, Fu J, Terpstra IR, Garcia JM, van den Ackerveken G, Snoek LB, et al. Regulatory network construction in Arabidopsis by using genome-wide gene expression quantitative trait loci. Proc Natl Acad Sci U S A. 2007;104: 1708–13. doi:10.1073/pnas.0610429104

35. van Leeuwen H, Kliebenstein DJ, West MAL, Kim K, van Poecke R, Katagiri F, et al. Natural variation among Arabidopsis thaliana accessions for transcriptome response to exogenous salicylic acid. Plant Cell. 2007;19: 2099–110. doi:10.1105/tpc.107.050641

36. West MAL, Kim K, Kliebenstein DJ, van Leeuwen H, Michelmore RW, Doerge RW, et al. Global eQTL mapping reveals the complex genetic architecture of transcript-level variation in Arabidopsis. Genetics. 2007;175: 1441–50. doi:10.1534/genetics.106.064972

37. Zhang X, Cal AJ, Borevitz JO. Genetic architecture of regulatory variation in Arabidopsis thaliana. Genome Res. 2011;21: 725–733. doi:10.1101/gr.115337.110

38. Lowry DB, Logan TL, Santuari L, Hardtke CS, Richards JH, DeRose-Wilson LJ, et al. Expression quantitative trait locus mapping across water availability environments reveals contrasting associations with genomic features in Arabidopsis. Plant Cell. 2013;25: 3266–79. doi:10.1105/tpc.113.115352

39. Swanson-Wagner R a, DeCook R, Jia Y, Bancroft T, Ji T, Zhao X, et al. Paternal dominance of trans-eQTL influences gene expression patterns in maize hybrids. Science. 2009;326: 1118–1120. doi: 10.1126/science.1178294

40. Holloway B, Luck S, Beatty M, Rafalski J-A, Li B. Genome-wide expression quantitative trait loci (eQTL) analysis in maize. BMC Genomics. 2011;12: 336. doi:10.1186/1471-2164-12-336

41. Fu J, Cheng Y, Linghu J, Yang X, Kang L, Zhang Z, et al. RNA sequencing reveals the complex regulatory network in the maize kernel. Nat Commun. 2013;4: 2832. doi:10.1038/ncomms3832

42. Wang J, Yu H, Xie W, Xing Y, Yu S, Xu C, et al. A global analysis of QTLs for expression variations in rice shoots at the early seedling stage. Plant J. 2010;63: 1063–1074. doi:10.1111/j.1365-313X.2010.04303.x

43. Wang J, Yu H, Weng X, Xie W, Xu C, Li X, et al. An expression quantitative trait loci-guided co-expression analysis for constructing regulatory network using a rice recombinant inbred line population. J Exp Bot. 2014;65: 1069–1079. doi:10.1093/jxb/ert464

44. Kirst M, Basten CJ, Myburg AA, Zeng ZB, Sederoff RR. Genetic Architecture of Transcript-Level Variation in Differentiating Xylem of a Eucalyptus Hybrid. Genetics. 2005;169: 2295–2303. Available: http://www.genetics.org/cgi/content/abstract/169/4/2295

45. Drost DR, Benedict CI, Berg A, Novaes E, Novaes CRDB, Yu Q, et al. Diversification in the genetic architecture of gene expression and transcriptional networks in organ differentiation of Populus. Proc Natl Acad Sci U S A. 2010;107: 8492–7. doi:10.1073/pnas.0914709107

46. Kullan AR, van Dyk MM, Hefer C a, Jones N, Kanzler A, Myburg A a. Genetic dissection of growth, wood basic density and gene expression in interspecific backcrosses of Eucalyptus grandis and E. urophylla. BMC Genet. 2012;13: 60. doi:10.1186/1471-2156-13-60

47. Brem RB, Kruglyak L. The landscape of genetic complexity across 5,700 gene expression traits in yeast. Proc Natl Acad Sci U S A. 2005;102: 1572–7. doi:10.1073/pnas.0408709102

48. Hughes KA, Ayroles JF, Reedy MM, Drnevich JM, Rowe KC, Ruedi EA, et al. Segregating Variation in the Transcriptome: Cis Regulation and Additivity of Effects. Genetics. 2006;173: 1347–1355. doi:10.1534/genetics.105.051474

49. Meiklejohn CD, Parsch J, Ranz JM, Hartl DL. Rapid evolution of male-biased gene expression in Drosophila. Proc Natl Acad Sci U S A. 2003;100: 9894–9 doi:10.1073/pnas.1630690100

50. Potokina E, Druka A, Luo Z, Wise R, Waugh R, Kearsey M. Gene expression quantitative trait locus analysis of 16 000 barley genes reveals a complex pattern of genome-wide transcriptional regulation. Plant J. 2008;53: 90–101. doi:10.1111/j.1365-313X.2007.03315.x

51. Whitehead A, Crawford DL. Variation within and among species in gene expression: raw material for evolution. Mol Ecol. 2006;15: 1197–1211. doi:10.1111/j.1365-294X.2006.02868.x

52. Leder EH, McCairns RJS, Leinonen T, Cano JM, Viitaniemi HM, Nikinmaa M, et al. The Evolution and Adaptive Potential of Transcriptional Variation in Sticklebacks--Signatures of Selection and Widespread Heritability. Mol Biol Evol. 2015;32: 674–689. doi:10.1093/molbev/msu328

53. Mäkinen H, Papakostas S, Vøllestad LA, Leder EH, Primmer CR. Plastic and evolutionary gene expression responses are correlated in European grayling (Thymallus thymallus) subpopulations adapted to different thermal environments. J Hered. 2016;107: 82–89. doi:10.1093/jhered/esv069

54. Kohn MH, Shapiro J, Wu C-I. Decoupled differentiation of gene expression and coding sequence among Drosophila populations. Genes Genet Syst. 2008;83: 265–273. doi:10.1266/ggs.83.265

55. Holloway AK, Lawniczak MKN, Mezey JG, Begun DJ, Jones CD. Adaptive gene expression divergence inferred from population genomics. PLoS Genet. 2007;3: 2007–2013. doi:10.1371/journal.pgen.0030187

56. Leinonen T, McCairns RJS, O’Hara RB, Merilä J. *Q*_ST_–*F*_ST_ comparisons: evolutionary and ecological insights from genomic heterogeneity. Nat Rev Genet. 2013;14: 179–90. doi:10.1038/nrg3395

57. Nourmohammad A, Rambeau J, Held T, Berg J, Lassig M. Pervasive adaptation of gene expression in Drosophila. arXiv. 2015; Available: http://arxiv.org/abs/1502.06406

58. MacNeil LT, Walhout AJM. Gene regulatory networks and the role of robustness and stochasticity in the control of gene expression. Genome Res. 2011;21: 645–657. doi:10.1101/gr.097378.109

59. Whitacre JM, Bender A. Networked buffering: a basic mechanism for distributed robustness in complex adaptive systems. Theor Biol Med Model. 2010;7: 20. doi:10.1186/1742-4682-7-20

60. Whitacre JM. Biological robustness: Paradigms, mechanisms, systems principles. Front Genet. 2012;3: 1–15. doi:10.3389/fgene.2012.00067

61. Jansson S, Douglas CJ. Populus: A Model System for Plant Biology. Annu Rev Plant Biol. 2007;58: 435–458. doi:10.1146/annurev.arplant.58.032806.103956

62. Wang J, Street NR, Scofield DG, Ingvarsson PK. Variation in linked selection and recombination drive genomic divergence during allopatric speciation of European and American aspens. Mol Biol Evol. 2016; msw051. doi:10.1093/molbev/msw051

63. Wang J, Street NR, Scofield DG, Ingvarsson PK. Natural Selection and Recombination Rate Variation Shape Nucleotide Polymorphism Across the Genomes of Three Related Populus Species. Genetics. 2016;202: 1185–1200. doi:10.1534/genetics.115.183152

64. Sundell D, Mannapperuma C, Netotea S, Delhomme N, Lin Y-C, Sjödin A, et al. The Plant Genome Integrative Explorer Resource: PlantGenIE.org. New Phytol. 2015;208: 1149–1156. doi:10.1111/nph.13557

65. ftp://plantgenie.org/Data/PopGenIE/Populus_tremula/v1.1/VCF/SwAsp_94samples.filter.maf005.beagle.maf.recode.vcf.gz.

66. Luquez V, Hall D, Albrectsen BR, Karlsson J, Ingvarsson P, Jansson S. Natural phenological variation in aspen (Populus tremula): the SwAsp collection. Tree Genet Genomes. 2008;4: 279–292. doi:10.1007/s11295-007-0108-y

67. Ingvarsson PK. Nucleotide polymorphism and linkage disequilibrium within and among natural populations of European aspen (Populus tremula L., Salicaceae). Genetics. 2005;169: 945–53. doi:10.1534/genetics.104.034959

68. Hall D, Luquez V, Garcia VM, St Onge KR, Jansson S, Ingvarsson PK. Adaptive population differentiation in phenology across a latitudinal gradient in European aspen (Populus tremula, L.): a comparison of neutral markers, candidate genes and phenotypic traits. Evolution. 2007;61: 2849–60. doi:10.1111/j.1558-5646.2007.00230.x

69. Wright FA, Sullivan PF, Brooks AI, Zou F, Sun W, Xia K, et al. Heritability and genomics of gene expression in peripheral blood. Nat Genet. 2014;46: 430–7. doi:10.1038/ng.2951

70. Yang S, Liu Y, Jiang N, Chen J, Leach L, Luo Z, et al. Genome-wide eQTLs and heritability for gene expression traits in unrelated individuals. BMC Genomics. 2014;15: 13. doi:10.1186/1471-2164-15-13

71. Zan Y, Shen X, Forsberg SKG, Carlborg Ö. Genetic regulation of transcriptional variation in wild-collected Arabidopsis thaliana accessions. bioRxiv. 2016; Available: http://biorxiv.org/content/early/2016/01/18/037077.abstract

72. Schmitz RJ, Schultz MD, Urich MA, Nery JR, Pelizzola M, Libiger O, et al. Patterns of population epigenomic diversity. Nature. 2013;495: 193–8. doi:10.1038/nature11968

73. Dubin MJ, Zhang P, Meng D, Remigereau M-S, Osborne EJ, Paolo Casale F, et al. DNA methylation in Arabidopsis has a genetic basis and shows evidence of local adaptation. Elife. 2015;4: e05255. doi:10.7554/eLife.05255

74. Robinson KM, Delhomme N, Mähler N, Schiffthaler B, Onskog J, Albrectsen BR, et al. Populus tremula (European aspen) shows no evidence of sexual dimorphism. BMC Plant Biol. 2014;14: 276. doi:10.1186/s12870-014-0276-5

75. Hyun MK, Ye C, Eskin E. Accurate discovery of expression quantitative trait loci under confounding from spurious and genuine regulatory hotspots. Genetics. 2008;180: 1909–1925. doi:10.1534/genetics.108.094201

76. Pickrell JK, Marioni JC, Pai AA, Degner JF, Engelhardt BE, Nkadori E, et al. Understanding mechanisms underlying human gene expression variation with RNA sequencing. Nature. 2010;464: 768–72. doi:10.1038/nature08872

77. Mostafavi S, Battle A, Zhu X, Urban AE, Levinson D, Montgomery SB, et al. Normalizing RNA-sequencing data by modeling hidden covariates with prior knowledge. PLoS One. 2013;8: e68141. doi:10.1371/journal.pone.0068141

78. Battle A, Mostafavi S, Zhu X, Potash JB, Weissman MM, McCormick C, et al. Characterizing the genetic basis of transcriptome diversity through RNA-sequencing of 922 individuals. Genome Res. 2014;24: 14–24. doi:10.1101/gr.155192.113

79. Massouras A, Waszak SM, Albarca-Aguilera M, Hens K, Holcombe W, Ayroles JF, et al. Genomic variation and its impact on gene expression in Drosophila melanogaster. PLoS Genet. 2012;8: e1003055. doi:10.1371/journal.pgen.1003055

80. Barabási A-L, Oltvai ZN. Network biology: understanding the cell’s functional organization. Nat Rev Genet. 2004;5: 101–13. doi:10.1038/nrg1272

81. Sjödin A, Street NR, Sandberg G, Gustafsson P, Jansson S. The Populus Genome Integrative Explorer (PopGenIE): a new resource for exploring the Populus genome. New Phytol. 2009;182: 1013–1025. doi:10.1111/j.1469-8137.2009.02807.x

82. Dai X, Hu Q, Cai Q, Feng K, Ye N, Tuskan GA, et al. The willow genome and divergent evolution from poplar after the common genome duplication. Cell Res. Shanghai Institutes for Biological Sciences, Chinese Academy of Sciences; 2014;24: 1274–1277. doi:10.1038/cr.2014.83

83. Tung J, Zhou X, Alberts SC, Stephens M, Gilad Y. The genetic architecture of gene expression levels in wild baboons. Elife. 2015;4: 1–22. doi:10.7554/eLife.04729

84. Wagner A. Genotype networks shed light on evolutionary constraints. Trends Ecol Evol. Elsevier Ltd; 2011;26: 577–584. doi:10.1016/j.tree.2011.07.001

85. Alvarez-Ponce D, Aguadé M, Rozas J. Network-level molecular evolutionary analysis of the insulin / TOR signal transduction pathway across 12 Drosophila genomes. Genome Res. 2009;19: 234–42. doi:10.1101/gr.084038.108.234

86. Zhang W, Landback P, Gschwend AR, Shen B, Long M. New genes drive the evolution of gene interaction networks in the human and mouse genomes. Genome Biol. 2015;16: 202. doi:10.1186/s13059-015-0772-4

87. Popadin KY, Gutierrez-Arcelus M, Lappalainen T, Buil A, Steinberg J, Nikolaev SI, et al. Gene age predicts the strength of purifying selection acting on gene expression variation in humans. Am J Hum Genet. 2014;95: 660–74. doi:10.1016/j.ajhg.2014.11.003

88. Zhang J, Yang J-R. Determinants of the rate of protein sequence evolution. Nat Rev Genet. Nature Publishing Group; 2015;16: 409–420. doi:10.1038/nrg3950

89. Albert R, Barabasi AL. Statistical mechanics of complex networks. Rev Mod Phys. 2002;74: 47–97. doi:10.1088/1478-3967/1/3/006

90. Choi J, Shooshtari P, Samocha KE, Daly MJ, Cotsapas C. Network Analysis of Genome-Wide Selective Constraint Reveals a Gene Network Active in Early Fetal Brain Intolerant of Mutation. PLoS Genet. 2016;12: 1–16. doi:10.1371/journal.pgen.1006121

91. Battle A, Mostafavi S, Zhu X, Potash JB, Weissman MM, McCormick C, et al. Characterizing the genetic basis of transcriptome diversity through RNA-sequencing of 922 individuals. Genome Res. 2013; doi:10.1101/gr.155192.113

92. Papakostas S, Vøllestad LA, Bruneaux M, Aykanat T, Vanoverbeke J, Ning M, et al. Gene pleiotropy constrains gene expression changes in fish adapted to different thermal conditions. Nat Commun. 2014;5: 4071. doi:10.1038/ncomms5071

93. Robinson KM, Ingvarsson PK, Jansson S, Albrectsen BR. Genetic variation in functional traits influences arthropod community composition in aspen (Populus tremula L.). PLoS One. 2012;7: e37679. doi:10.1371/journal.pone.0037679

94. Chang S, Puryear J, Cairney J. A simple and efficient method for isolating RNA from pine trees. Plant Mol Biol Report. 1993;11: 113–116. doi:10.1007/BF02670468

95. Leinonen R, Akhtar R, Birney E, Bower L, Cerdeno-Tárraga A, Cheng Y, et al. The European Nucleotide Archive. Nucleic Acids Res. 2011;39: D28–31. doi:10.1093/nar/gkq967

96. Delhomme N, Mähler N, Schiffthaler B, Sundell D, Mannepperuma C, Hvidsten TR, et al. Guidelines for RNA-Seq data analysis. Epigenesys. 2014; Available: http://www.epigenesys.eu/en/protocols/bio-informatics/1283-guidelines-for-rna-seq-data-analysis

97. Bolger AM, Lohse M, Usadel B. Trimmomatic: a flexible trimmer for Illumina sequence data. Bioinformatics. 2014;30: 2114–2120. doi:10.1093/bioinformatics/btu170

98. Kopylova E, Noé L, Touzet H. SortMeRNA: Fast and accurate filtering of ribosomal RNAs in metatranscriptomic data. Bioinformatics. 2012;28: 3211–3217. doi:10.1093/bioinformatics/bts611

99. Dobin A, Davis CA, Schlesinger F, Drenkow J, Zaleski C, Jha S, et al. STAR: ultrafast universal RNA-seq aligner. Bioinformatics. 2012;29: 15–21. doi:10.1093/bioinformatics/bts635

100. Anders S, Pyl PT, Huber W. HTSeq - A Python framework to work with high-throughput sequencing data. Bioinformatics. 2014;31: 166–169. doi:10.1093/bioinformatics/btu638

101. Andrews S. FastQC: A quality control tool for high throughput sequence data [Internet]. 2016 [cited 27 Sep 2016]. Available: http://www.bioinformatics.bbsrc.ac.uk/projects/fastqc/

102. Love MI, Huber W, Anders S. Moderated estimation of fold change and dispersion for RNA-seq data with DESeq2. Genome Biol. 2014;15: 550. doi:10.1186/s13059-014-0550-8

103. Dohm MR. Repeatability estimates do not always set an upper limit to heritability. Funct Ecol. 2002;16: 273–280. doi:10.1046/j.1365-2435.2002.00621.x

104. Kruijer W, Boer MP, Malosetti M, Flood PJ, Engel B, Kooke R, et al. Marker-based estimation of heritability in immortal populations. Genetics. 2015;199: 379–398. doi:10.1534/genetics.114.167916

105. Spitze K. Population structure in Daphnia obtusa: quantitative genetic and allozymic variation. Genetics. 1993;135: 367–74. Available: http://www.genetics.org/content/135/2/367.short

106. Bates D, Mächler M, Bolker B, Walker S. Fitting Linear Mixed-Effects Models Using lme4. J Stat Softw. 2015;67: 1–48. doi:10.18637/jss.v067.i01

107. Manier MK, Palumbi SR. Intraspecific divergence in sperm morphology of the green sea urchin, Strongylocentrotus droebachiensis: implications for selection in broadcast spawners. BMC Evol Biol. 2008;8: 283. doi:10.1186/1471-2148-8-283

108. Listgarten J, Kadie C, Schadt EE, Heckerman D. Correction for hidden confounders in the genetic analysis of gene expression. Proc Natl Acad Sci U S A. 2010;107: 16465–70. doi:10.1073/pnas.1002425107

109. Parts L, Stegle O, Winn J, Durbin R. Joint genetic analysis of gene expression data with inferred cellular phenotypes. PLoS Genet. 2011;7: 1–10. doi:10.1371/journal.pgen.1001276

110. Shabalin AA. Matrix eQTL: Ultra fast eQTL analysis via large matrix operations. Bioinformatics. 2012;28: 1353–1358. doi:10.1093/bioinformatics/bts163

111. Purcell S, Neale B, Todd-Brown K, Thomas L, Ferreira MA, Bender D, et al. PLINK: a tool set for whole-genome association and population-based linkage analyses. Am J Hum Genet. 2007;81: 559–575. doi:10.1086/519795

112. Storey J. qvalue: Q-value estimation for false discovery rate control [Internet]. 2015. Available: http://github.com/jdstorey/qvalue

113. Langfelder P, Horvath S. WGCNA: an R package for weighted correlation network analysis. BMC Bioinformatics. 2008;9: 559. doi:10.1186/1471-2105-9-559

114. ftp://plantgenie.org/Data/PopGenIE/Populus_tremula/v1.1/Annotation/DoubleMCL-tab-annotation.txt.gz.

115. Jin J, Zhang H, Kong L, Gao G, Luo J. PlantTFDB 3.0: A portal for the functional and evolutionary study of plant transcription factors. Nucleic Acids Res. 2014;42: 1182–1187. doi:10.1093/nar/gkt1016

116. Tajima F. Statistical method for testing the neutral mutation hypothesis by DNA polymorphism. Genetics. 1989;123: 585–595. doi:PMC1203831

117. Korneliussen T, Albrechtsen A, Nielsen R. ANGSD: Analysis of Next Generation Sequencing Data. BMC Bioinformatics. 2014;15: 356. doi:10.1186/s12859-014-0356-4

118. Zhang C, Wang J, Long M, Fan C. GKaKs: The pipeline for genome-level Ka/Ks calculation. Bioinformatics. 2013;29: 645–646. doi:10.1093/bioinformatics/btt009

119. Yang Z. PAML 4: Phylogenetic Analysis by Maximum Likelihood. Mol Biol Evol. 2007;24: 1586–1591. doi:10.1093/molbev/msm088

